# Gene therapies alleviate absence epilepsy associated with *Scn2a* deficiency in DBA/2J mice

**DOI:** 10.1101/2025.06.03.657652

**Authors:** Zaiyang Zhang, Jingliang Zhang, Xiaoling Chen, Brody A. Deming, Shivam Kant, Purba Mandal, Harish Kothandaraman, Phillip J. SanMiguel, Manasi S. Halurkar, Akila D. Abeyaratna, Morgan J. Robinson, Yuanrui Zhao, Yuliia Vitko, Ronald P. Gaykema, Chongli Yuan, Nadia A. Lanman, Matthew T. Tegtmeyer, Dan Wang, Guangping Gao, Riyi Shi, Edward Perez-Reyes, Yang Yang

## Abstract

Mutations in the voltage-gated sodium channel gene *SCN2A*, which encodes the Na_V_1.2 channel, cause severe epileptic seizures. Patients with *SCN2A* loss-of-function (LoF) mutations, such as protein-truncating mutations, often experience later-onset and drug-resistant epilepsy, highlighting an urgent unmet clinical need for new therapies. We previously developed a gene-trap *Scn2a* (*Scn2a^gt/gt^*) mouse model with a global Na_V_1.2 reduction in the widely used C57BL/6N (B6) strain. Although these mice display multiple behavioral abnormalities, EEG recordings indicated only mild epileptiform discharges, possibly attributable to the seizure-resistant characteristics associated with the B6 strain. To enhance the epileptic phenotype, we derived congenic *Scn2a^gt/gt^* mice in the seizure-susceptible DBA/2J (D2J) strain. Notably, we found that these mice exhibit prominent spontaneous absence seizures, marked by both short and long spike-wave discharges (SWDs). Restoring Na_V_1.2 expression in adult mice substantially reduced their SWDs, suggesting the possibility of *SCN2A* gene replacement therapy during adulthood. RNA sequencing revealed significant alterations in gene expression in the *Scn2a^gt/gt^* mice, in particular a broad downregulation of voltage-gated potassium channel (K_V_) genes, including K_V_1.1. The reduction of K_V_1.1 expression was further validated in human cerebral organoids with *SCN2A* deficiency, highlighting K_V_1.1 as a promising therapeutic target for refractory seizures associated with *SCN2A* dysfunction. Importantly, delivery of exogenous human K_V_1.1 expression via adeno-associated virus (AAV) in D2J *Scn2a^gt/gt^* mice substantially reduced absence seizures. Together, these findings underscore the influence of mouse strain on seizure severity and highlight the potential of targeted gene therapies for treating *SCN2A* deficiency-related epilepsies.

**Graphical abstract:** 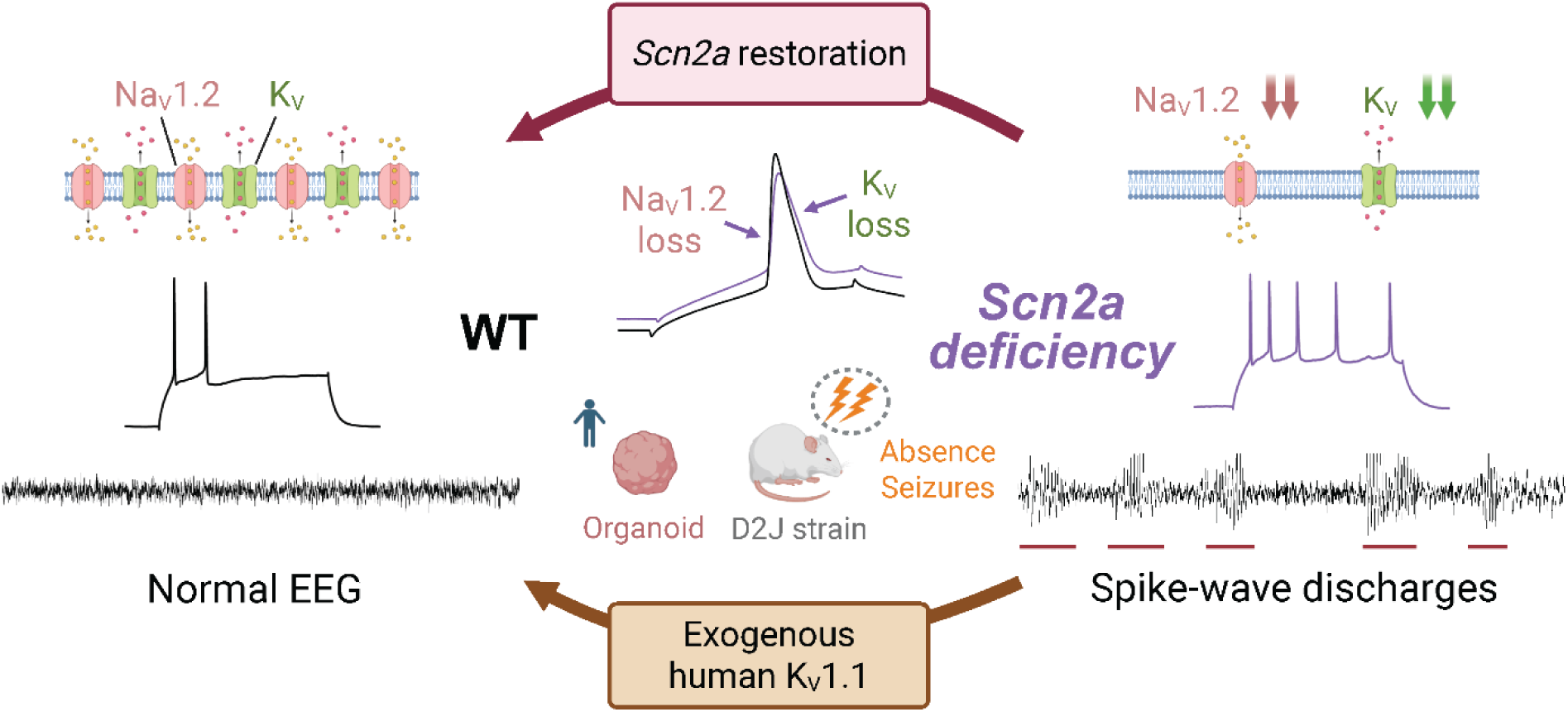

*In brief:* *Scn2a* deficiency leads to absence seizures in D2J mice and neuronal hyperexcitability with compensatory K_V_ reduction; restoring Na_V_1.2 or introducing human K_V_1.1 reduces seizure burden.

**Highlights:**

1. *Scn2a* deficiency induces robust absence seizures in the DBA/2J but not the C57BL/6N strain.
2. Cortical neurons in adult DBA/2J mice with *Scn2a* deficiency exhibit intrinsic hyperexcitability.
3. Severe *Scn2a* deficiency leads to downregulation of multiple potassium channel genes.
4. Genetic restoration of Na_V_1.2 expression alleviates spike-wave discharges (SWDs).
5. AAV-mediated human K_V_1.1 delivery substantially reduced absence seizures, demonstrating the therapeutic potential of targeted gene therapy.

## Introduction

Epilepsy is a chronic disease marked by recurrent unprovoked seizures. With at least 50 million epileptic patients worldwide, it is one of the most prevalent neurological disorders (1). More than 70% of epilepsy cases have a genetic component (2), and *de novo* single gene variants account for 30%–50% of developmental epileptic encephalopathies (DEEs) (3). Recent advances in diagnostic sequencing have identified *SCN2A* as one of the top three monogenic variants in patients with developmental epilepsy, underscoring its pivotal role in epileptogenesis (4). The *SCN2A* gene encodes for the alpha subunit of the voltage-gated sodium ion channel 1.2 (Na_V_1.2), and mutations can occur in any part of its domains, leading to a spectrum of complex cellular and behavioral phenotypes (5, 6). These mutations can be roughly categorized into gain-of-function (GoF) or loss-of-function (LoF), depending on the biophysical properties affected (7). Patients with *SCN2A* GoF mutations typically present with early-onset epilepsy and respond well to antiepileptic drugs (AEDs), whereas those with LoF mutations often experience later-onset and drug-resistant epilepsy (8). Individuals with *SCN2A* LoF mutations display various types of seizures, including absence epilepsy, a type of generalized seizure marked by abrupt, brief lapses in consciousness and spike-and-wave discharges (SWDs) in EEG (5, 9, 10). A recent comprehensive clinical study conducted functional phenotyping for mutations from a large cohort of *SCN2A* patients, in which 71.6% of the variants were categorized into LoF (5). Moreover, among the complete LoF patients (i.e., truncation mutation), as much as 65% presented seizures, indicating that *SCN2A* LoF-related epilepsy affects a sizable patient population (5).

To model *Scn2a* complete LoF mutations, the generation of *Scn2a* knockout (KO) mice had been previously attempted (11). However, heterozygous knockout renders modest seizure-like phenotypes (12) and complete germline deletion of *Scn2a* in mice led to perinatal mortality, likely due to its indispensable role in action potential regulation during neurodevelopment (11). To tackle this obstacle, our lab developed a *Scn2a* gene-trap (*Scn2a^gt/gt^*) mouse model in the widely used C57BL/6N (B6) strain (13) that exhibits a significant global reduction in Na_V_1.2 expression (∼30% expression of the wild-type (WT) mice), while remaining viable into adulthood (14, 15). While these mice display multiple behavioral abnormalities, seizure-related EEG phenotypes were mild. We hypothesized that the lack of strong seizure-related phenotypes could be partially due to the inherent seizure-resistant nature associated with the B6 strains. Strain-dependent seizure severity has been validated in a variety of epilepsy models, including post-traumatic epilepsy (16), chemical kindling (17, 18), electrical stimulation (19), and genetic epilepsy (20–22). In contrast to the B6 strains, the DBA/2J (D2J) strain is recognized as one of the most seizure-susceptible strains (17, 23). Thus, we enhanced the seizure phenotypes by crossing *Scn2a^gt/gt^* mice into the D2J strain and then used it as a preclinical disease model to evaluate potential disease intervention strategies. While current AEDs are small molecules, the development of gene therapy holds great promise for drug-resistant seizures with known genetic causes. Gene therapies have been tested in the *Scn1a*-deficient mouse model of Dravet syndrome as well as other animal models of monogenic epilepsy, achieving encouraging effects in reducing seizure burdens (24, 25). However, the effect of genetic-based approaches on seizure-related phenotypes in *Scn2a*-deficient mice has not been reported.

In this study, we found that *Scn2a*-deficient mice in the D2J strain, rather than the B6 strain, display robust absence seizure phenotype characterized by repeated spike-wave discharges (SWDs). Partial restoration of Na_V_1.2 expression in adulthood alleviated their SWDs. Because Na_V_1.2 interacts with multiple functionally related proteins, severe Na_V_1.2 reduction causes widespread multichannel disturbances, with closely associated channels up- or down-regulated in compensation during neurodevelopment (26). Therefore, guided by altered neuronal electrophysiological properties and differential gene expression from bulk RNA sequencing, we identified robust compensatory downregulation of voltage-gated potassium channel genes as alternative targets. Considering AAV-K_V_1.1 has been demonstrated as a promising gene therapy in various epilepsy animal models (27, 28), we assessed the effect of exogenous human K_V_1.1 delivery in adulthood on D2J *Scn2a^gt/gt^* mice. Notably, we found that the absence seizures were significantly alleviated using this strategy. Our results demonstrated the utility of D2J *Scn2a^gt/gt^* mice as a disease model and shed light on future translational endeavors for treating *SCN2A-*related seizures using different genetic approaches.

## Results

### Generation of the congenic gene-trap Scn2a-deficient mice in the DBA/2J (D2J) strain

Our lab has previously generated *Scn2a*-deficient (*Scn2a^gt/gt^*) mice in the widely used C57BL/6N (B6) strain, which display multiple behavioral abnormalities, including social deficits, impaired innate behavior and disrupted circadian rhythm (14, 29, 30). However, these mice display mild seizure-related phenotypes (**Supplemental Figure 1**). To examine possible epileptiform discharges in the *Scn2a^gt/gt^* mice, we conducted one-week continuous video-EEG recordings. Prefabricated headmounts were used to capture cortical neuronal activities and simultaneous electromyography (EMG) signals, which indicate animal movement (**Figure 1**). We detected statistically significant but mild increased short spike-wave discharges (S-SWDs) in the B6 *Scn2a^gt/gt^* mice compared to their B6 wild-type (WT) littermates. These SWDs were detected in both anterior and posterior cortical electrodes, with the frontal cortex exhibiting a stronger signal. Therefore, we used the anterior electrode recordings for all the following SWD quantifications. In contrast, no SWD activity was detected in either recording electrode in the B6 WT mice (0 for B6-WT vs. 0.03 ± 0.01 per hour for B6-*Scn2a^gt/gt^*; **p < 0.01, Mann-Whitney U test) (**Supplemental Figure 1, A-B**). These data suggest that mice in the B6 genetic background might not be suitable for modeling *SCN2A* deficiency patients suffering from severe epilepsy.

**Figure 1.**
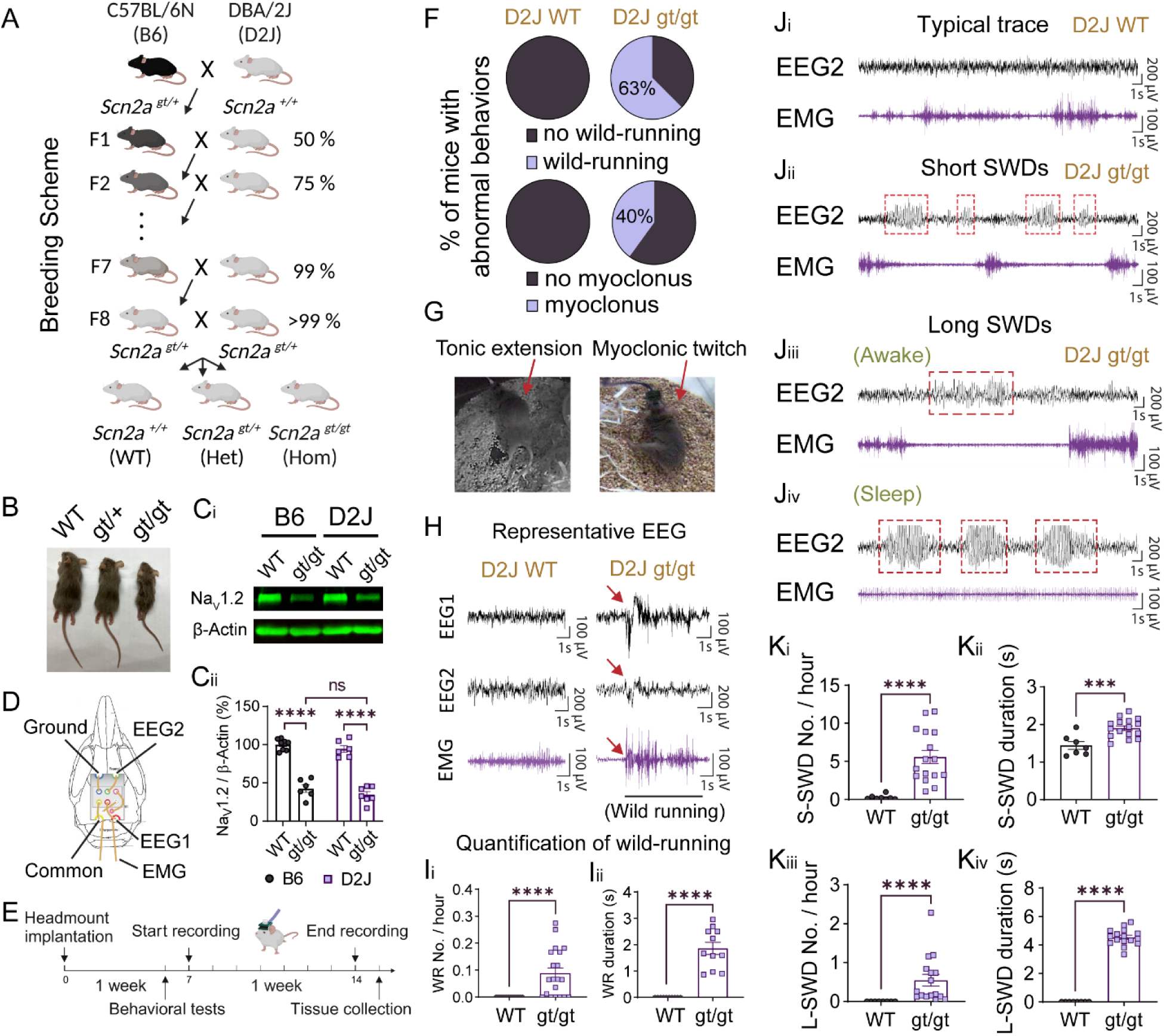
Severe deficiency of *Scn2a* results in spontaneous absence seizures and abnormal behaviors in DBA/2J mice. (A) The breeding scheme for generating the *Scn2a* gene-trap congenic mice in the DBA/2J (D2J) background from the C57BL/6N (B6) background. (B) D2J *Scn2a^gt/gt^* mice have smaller body sizes compared to D2J WT or D2J Het mice. (C) Western blot shows that the Na_V_1.2 protein expression level in the D2J *Scn2a^gt/gt^* mice is decreased to a similar level to the B6 *Scn2a^gt/gt^* mice compared to their corresponding WTs. Results were normalized to the expression of housekeeping protein β-actin. N = 6 for all four groups. (D) A schematic of the prefabricated 2EEG/1EMG headmount from the Pinnacle system. (E) Timeline of the EEG recording experiment. Mice were recovered in their home cage for at least a week before 1-week video-EEG recording. (F) Percentage of D2J *Scn2a^gt/gt^* mice with wild-running (WR) behavior or myoclonic twitches. (G) Video screenshots show epilepsy-related abnormal behaviors in two D2J *Scn2a^gt/gt^* mice. (H) Representative EEG1 (posterior) and EEG2 (anterior) signals for a typical WT mouse awake and walking and a *Scn2a^gt/gt^* D2J mouse during the wild-running episode. Red arrows indicate the start of the wild running event. (I) Quantification of the frequency and duration of the wild-running (WR) behavior for D2J WT and D2J *Scn2a^gt/gt^*. (J) Representative EEG2 and EMG traces of typical D2J WT mouse with no spike-wave discharges (SWD), D2J *Scn2a^gt/gt^* mice with short SWDs (S-SWDs), and D2J *Scn2a^gt/gt^* mice with long SWDs (L-SWDs) which last > 3.5 s during awake or asleep states. (K) Quantifications of the frequency and duration of short and long spike-wave discharges (SWD) in the D2J WT and *Scn2a^gt/gt^*. Data are presented as mean ± SEM. Statistical analyses: Two-way ANOVA: F (DFn, DFd) = F (1, 23) = 221.8 for the genotype factor and post hoc multiple comparisons with Tukey’s correction (B6 WT vs. *Scn2a^gt/gt^* and D2J WT vs. *Scn2a^gt/gt^*) (C). Mann-Whitney U test (Ii, Ki, Kiii, Kiv). Unpaired t-test (Kii). *p < 0.05; **p < 0.01; ***p < 0.001; ****p < 0.0001. Exact p values can be found in Table S1.

In patients with *SCN2A* truncating mutations, epilepsy manifests in only a subset, suggesting that individual genetic backgrounds may influence seizure susceptibility and epileptogenesis (5). Previous research indicates that C57BL/6 (B6) mice are more seizure-resistant, whereas strains such as DBA/2J (D2J) are more seizure-susceptible (17, 23). Therefore, we generated congenic *Scn2a* gene-trap transgenic mice in the D2J background by backcrossing the B6 *Scn2a^gt/+^* mice to inbred D2J WT mice for over eight generations (**Figure 1A**). The genomic background of the resulting D2J congenic mice was validated by the Giga Mouse Universal Genotyping Array (GigaMUGA), which showed 99.9% genome identity consistent with WT inbred D2J mice (**Supplemental Figure 1C**). We then crossed the D2J *Scn2a^gt/+^*offspring to obtain a colony of D2J *Scn2a^+/+^* (WT), *Scn2a^gt/+^*(heterozygotes), and *Scn2a^gt/gt^* (homozygotes) mice (**Figure 1A**). The D2J *Scn2a^gt/gt^* mice had a smaller body size than the D2J *Scn2a^gt/+^* or *Scn2a^+/+^* mice (**Figure 1B**), a feature consistent with *Scn2a^gt/gt^* mice in the B6 background as reported previously (14). Since we did not detect obvious behavioral or EEG abnormality in the *Scn2a^gt/+^*heterozygous mice (data now shown), all experiments in this study were done using homozygotes. Importantly, whole brain western blot showed that Na_V_1.2 protein in the D2J *Scn2a^gt/gt^* mice had a similarly low level of expression comparable to the B6 *Scn2a^gt/gt^*mice, which was around 34% of their corresponding WTs (**Figure 1C**) This result confirms that our congenic strain generation did not interfere with the gene-trap-induced *Scn2a* expression reduction. In summary, we established a *Scn2a*-deficient mouse model in the seizure-prone DBA/2A strain that survived through adulthood.

### D2J Scn2a^gt/gt^ mice have prominent absence seizures

To determine if the *Scn2a* deficient mice in the ‘seizure-prone’ DBA/2A strain show major epileptiform discharges in the cortex, we conducted continuous EEG recordings as previously. We found that D2J *Scn2a^gt/gt^* mice displayed robust SWDs (**Figure 1**). These SWDs appear in both the anterior and posterior cortical EEG electrodes, in line with the characteristic of absence seizure as a type of generalized seizure. Short SWDs (S-SWD, 0–3.5 s) were detected in both D2J *Scn2a^gt/gt^* and D2J WT mice, but with significantly higher frequency and longer duration in the D2J *Scn2a^gt/gt^* mice (**Figure 1, J_i_-J_ii_ and K_i_-K_ii_**): D2J *Scn2a^gt/gt^* mice had an average of 5.57 ± 0.86 S-SWDs per hour whereas D2J WT mice only had 0.24 ± 0.11 S-SWDs per hour; the average duration of S-SWD was 1.90 ± 0.06 s for D2J *Scn2a^gt/gt^* mice vs. 1.44 ± 0.10 s for D2J WT mice. Long SWDs (L-SWD, >3.5 s) were detected only in the D2J *Scn2a^gt/gt^* mice and rarely in D2J WT mice (**Figure 1, J_iii_-J_iv_ and K_iii_-K_iv_**). These L-SWDs are characterized by prolonged duration, high amplitudes, elevated EEG power, and extended animal behavioral arrest, which more closely resembles absence epilepsy (31, 32). In D2J *Scn2a^gt/gt^* mice, L-SWD occurred on average 0.54 ± 0.15 per hour with a duration of 4.54 ± 0.14 s, while no L-SWD was observed in naïve D2J WT mice (****p < 0.0001; Mann-Whitney U test). The SWDs in D2J *Scn2a^gt/gt^*mice showed much higher frequency and intensity than the weak SWDs occasionally occurring in the D2J WT mice, indicating that *Scn2a* reduction induces severe absence seizures.

To thoroughly validate the severity of seizure phenotypes in D2J versus B6 *Scn2a*-deficient mice, we performed a detailed EEG signal comparison analysis to detect SWDs, which are identified as 5–7 Hz (33). The SWD frequency and duration in B6 *Scn2a^gt/gt^* mice were much less than those in the D2J *Scn2a^gt/gt^* mice (S-SWD frequency: 0.03 ± 0.01 for B6-*Scn2a^gt/gt^* vs. 4.42 ± 1.05 per hour for D2J-*Scn2a^gt/gt^*; ***p < 0.001; S-SWD duration: 1.47 ± 0.14 s for B6-*Scn2a^gt/gt^*vs. 1.75 ± 0.05 s for D2J-*Scn2a^gt/gt^*) (**Supplemental Figure 1B**). The shape of short SWDs in the B6 *Scn2a^gt/gt^* mice also appeared more ‘immature’ (34), with lower duration and amplitude compared to the D2J *Scn2a^gt/gt^* mice (**Supplemental Figure 1A**). Additionally, long SWDs (>3.5 s) were not observed in the B6 *Scn2a^gt/gt^* mice (**Supplemental Figure 1, Biii and Bvi**). These results align with our hypothesis that *Scn2a^gt/gt^*mice in the D2J strain have much stronger epileptiform discharges than those in the B6 strain. Collectively, we demonstrated that the *Scn2a*-deficient mice in the D2J background exhibit severe absence-like seizures, recapitulating one of the seizure phenotypes observed in *SCN2A* LoF patients (5, 9, 10).

### Scn2a-deficient mice in D2J background display seizure-related abnormal behaviors

To inspect further *Scn2a* deficiency-related phenotype, we carefully studied continuous EEG-video recordings and discovered that the D2J *Scn2a^gt/gt^*mice exhibit unique, abnormal behaviors that are absent in the D2J WT mice. Besides SWDs, we observed that 63% of the D2J *Scn2a^gt/gt^* mice displayed unprovoked wild-running (WR) behavior, and around 40% demonstrated myoclonic twitches (**Figure 1, F-G**). WR has been reported in rodents with seizures (35). In D2J *Scn2a^gt/gt^* mice with WR behavior, the frequency of WR averaged 0.09 per hour (ranging from 0.01 to 0.27 per hour) with a duration averaging around 2 seconds for 1 week of EEG recording (**Figure 1I**). Importantly, this behavior is completely absent in the D2J and B6 WT mice (**Figure 1I**). During WR, only the posterior electrode showed high amplitude signals, possibly due to its close approximation to the nuchal EMG electrodes. In contrast, no epileptiform activity was detected in the anterior electrode (**Figure 1H**). This finding aligns with earlier studies indicating that WR behavior may arise from subcortical structures (35). Likewise, myoclonic jerks are a well-established seizure-associated phenotype and have been characterized in both human patients and genetic mouse models (8, 36). Only a subset of D2J *Scn2a^gt/gt^* mice exhibits these abnormal behaviors (**Figure 1F**), suggesting possible heterogeneous developmental deficits linked to low Na_V_1.2 levels. D2J *Scn2a^gt/gt^* mice were also more hyperactive in the open-field test (OFT): They traveled a significantly longer distance at a higher speed, which increased their crossings to the center zone (**Supplemental Figure 3C**).

### Absence seizures in the D2J Scn2a^gt/gt^ mice are more severe during NREM sleep and appear clustered

Seizures are associated with circadian rhythms (e.g., sleep-wake cycle) in both patients and mice (37, 38), with SWDs dependent on vigilance, occurring most frequently during passive behavioral states or slow-wave sleep (39). Interestingly, we found that most absence seizures happened when the D2J *Scn2a^gt/gt^* mice were asleep (i.e. NREM state): L-SWD frequency for D2J-*Scn2a^gt/gt^* was 0.20 ± 0.06 per hour during sleep vs. 0.05 ± 0.02 per hour during awake ([Sleep - Awake] *Scn2a^gt/gt^* *adjusted p = 0.02; two-way ANOVA) (**Supplemental Figure 2**). The duration of L-SWD showed no difference between sleep and awake states (5.00 ± 0.19 s during sleep vs. 4.73 ± 0.23 s during awake; adjusted p = 0.43), indicating that absence seizures occurred more frequently, but not longer, when the mice were asleep (**Supplemental Figure 2D**). While SWDs were particularly severe during sleep, they also occur in the awake state. Notably, a ‘neck twitch’ usually happened at the onset of a long SWD episode when the mice were awake (**Supplemental Figure 2B, red arrows**). This neck muscle contraction has also been reported in other animal models with absence seizures (40, 41). Disruption of sleep-wake regulation has been explored in our previous study for the C57BL/6 (B6) *Scn2a^gt/gt^*mice (29) and is in line with absence seizure patients (10, 39). Interestingly, in both D2J and B6 mice, there was a consistent decrease in EEG baseline amplitude (μV) for *Scn2a^gt/gt^* mice during the NREM state, whereas no difference was observed in the awake state (**Supplemental Figure 2,**

**G-J**).

When examining the pattern of SWDs in D2J *Scn2a^gt/gt^* mice during 24-hour analysis, we observed that these epileptiform events tend to cluster instead of being evenly distributed, a trait frequently reported in human seizure patients (42) (**Supplemental Figure 3A**). Additionally, repeated trains of short SWDs that spread across several minutes were also frequently observed in the D2J *Scn2a^gt/gt^* mice (**Supplemental Figure 3B**). Throughout these events, mice transitioned from normal activities (e.g., eating, walking) to sudden behavioral arrest, characterized by a hunched posture, whisker twitching, staring, and body tottering, similar to other rodent models of absence seizure (34, 43).

### EEG recordings in Scn2a-deficient mice reveal reduced absolute power and altered relative power frequency distributions

Analysis of the EEG power frequency distribution allows us to identify seizure-associated oscillatory patterns and overall brain state alterations from the spectral profile. To characterize the overall EEG power frequency domain, we conducted power spectral analysis using the fast Fourier transform (FFT), which yielded measurements of power intensity per bandwidth, represented in units of Volt^2^/Hz (44). D2J *Scn2a^gt/gt^* mice showed robust power spectra differences compared to the D2J WT mice. D2J *Scn2a^gt/gt^* mice exhibit significantly reduced absolute power, particularly in the alpha, beta, and delta bands, as shown in the power spectral heatmap (**Figure 2, A and Ci-Cii**). The decrease was more pronounced during the light-on period, where quantification shows that total absolute power was significantly reduced (**Figure 2Di**). A similar overall absolute power reduction was observed in B6 *Scn2a^gt/gt^* mice compared to B6 WT mice, suggesting a conserved phenotype across the two mouse strains (**Supplemental Figure 1, Di-Dii**). Reduction in absolute power partially resembles EEG dysmaturity observed in humans, a feature of neurodevelopmental delay (45) and has been previously reported in *SCN1A*- and *SCN2A*-related epilepsies (46, 47).

**Figure 2.**
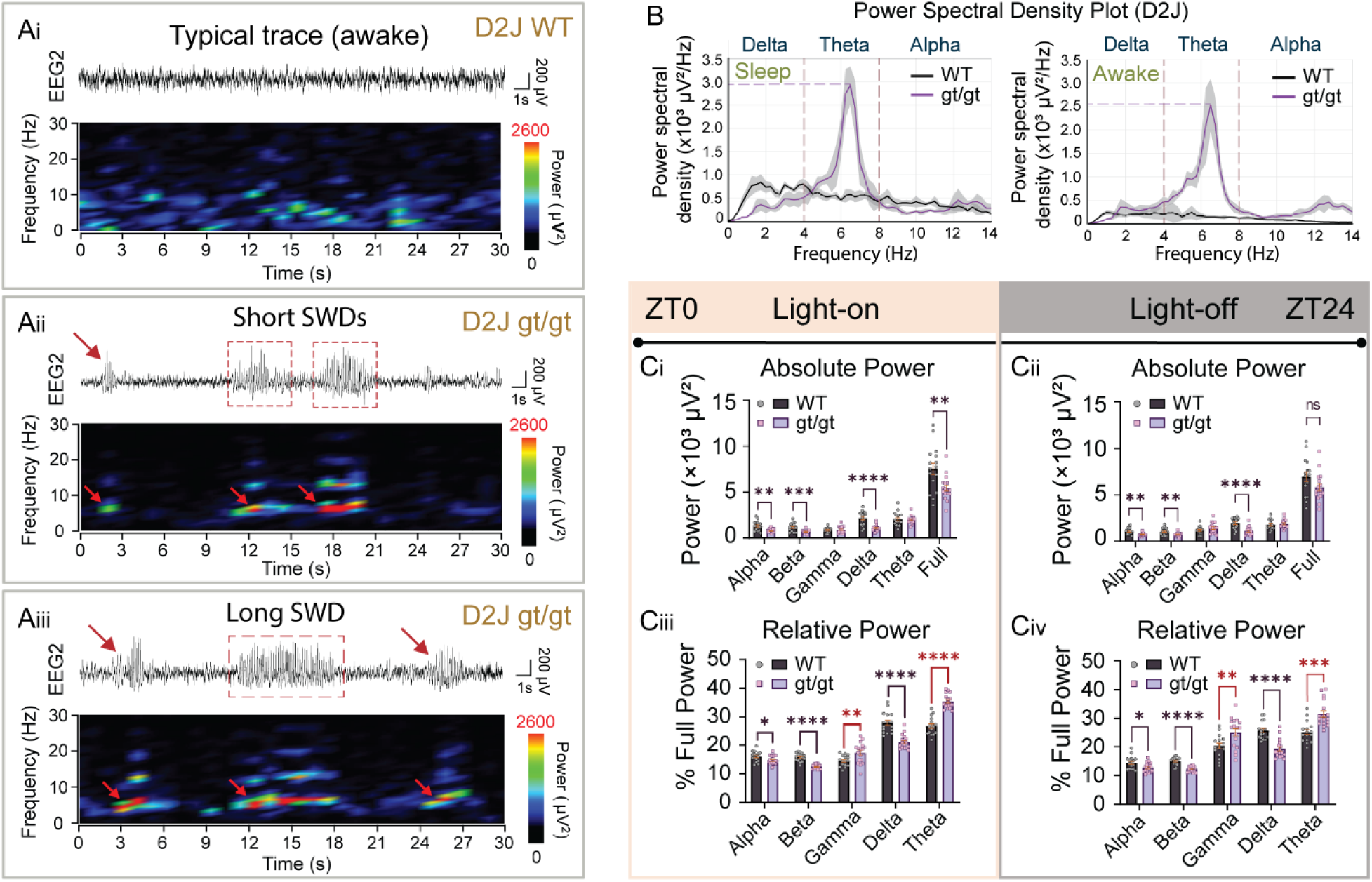
D2J *Scn2a^gt/gt^*mice show an overall reduction of absolute power with an increase in relative gamma and theta power. (A) Example EEG traces and corresponding power spectral density heatmaps. (Ai) A typical trace in a D2J WT mouse. (Aii) Two consecutive short spike-wave discharges (S-SWDs) in a D2J *Scn2a^gt/gt^* mouse. (Aiii) A long SWD accompanied by two S-SWDs (indicated by red arrows) in a D2J *Scn2a^gt/gt^* mouse. The animals were all in the awake state in these three examples. Note that the baseline EEG voltage is higher in the D2J WT than the D2J *Scn2a^gt/gt^* across all animals, according to the higher absolute power observed in the D2J WT mice. (B) Power spectral density plot showing examples of different frequency distributions for D2J WT vs. D2J *Scn2a^gt/gt^* mice during the light-on sleep (left) vs. light-off awake (right) states. Note that *Scn2a^gt/gt^* mice have high power density in the theta band (4–8Hz), corresponding to the frequency range of SWDs in mice. The maximum power of the theta band is higher during sleep compared to awake. (C) Quantification of absolute and relative power distribution of D2J WT and D2J *Scn2a^gt/gt^* mice over 1 week recording. Relative power was calculated by dividing the individual frequency band by the full power of that animal. (Ci-Cii) D2J *Scn2a^gt/gt^* mice have significantly lower absolute power compared to the D2J WT mice, especially in the alpha, beta, and delta frequency bands. (Ciii-Civ) D2J *Scn2a^gt/gt^*mice have lower relative power in the alpha, beta, and delta bands, but elevation of % power in the gamma and theta bands. Power spectra were calculated for light-on and light-off periods separately by Fast Fourier Transform (FFT) with Hann (cosine-bell) data window set using an epoch of 10 s based on the following frequency bands: full (0–100 Hz), delta (0.5–4 Hz), theta (4–8 Hz), alpha (8–13 Hz), beta (13–30 Hz), and gamma (30–100 Hz). The relative power was calculated by dividing each power band with the full power (0–100 Hz). Data are presented as mean ± SEM. Statistical analyses: Two-way ANOVA: F (DFn, DFd) = F (1, 186) = 29.01 ****p < 0.0001 for the [genotype] factor; Multiple t-test (C1). *p < 0.05; **p < 0.01; ***p < 0.001; ****p < 0.0001. Exact p values can be found in Supplemental Table 1.

We further calculated the relative power spectra by normalizing the specific power frequency band to the total power in each mouse. Consistent with the trend in absolute power, D2J *Scn2a^gt/gt^* mice displayed significantly lower relative power in the alpha, beta, and delta frequency bands. Similarly, in the B6 *Scn2a^gt/gt^* mice, there was a significant delta band decrease during the light-on period and an alpha band decrease during the light-off period (**Supplemental Figure 1, Diii-Div**). The delta frequency range (0.5–4 Hz) has been associated with slow wave (NREM) sleep (48). Interestingly, we discovered a lower EEG voltage during the NREM state in *Scn2a^gt/gt^* mice (**Supplemental Figure 2H**), which aligns with the significant reduction in EEG delta power. Studies on absence epilepsy suggest that the main frequency component of cortical SWDs lies within the theta band (49). Consistently, the relative power of the theta band was significantly elevated in D2J *Scn2a^gt/gt^* mice, and the increase was more prominent during the light-on period (**Figure 2, B and Ciii-Civ**, highlighted in red). Furthermore, as demonstrated in **Figure 2, Aii-Aiii**, during D2J *Scn2a^gt/gt^* mice SWD events, the power density of the theta band (4–8 Hz) was significantly elevated (highlighted by red arrows). In contrast, although B6 *Scn2a^gt/gt^* mice didn’t show a statistically significant difference in relative theta power compared to B6 WT mice (likely due to their infrequent SWD activity), a trend of increase was observed (**Supplemental Figure 1, Diii-Div**). Moreover, we found an elevation in relative gamma power, denoting high-frequency oscillations in the D2J *Scn2a^gt/gt^* mice (**Figure 2, Ciii-Civ)**. Gamma oscillations, interestingly, are hypothesized to be related to an increase in the action potential firing rate or hypersynchrony by assemblies of neurons (50). Therefore, the increase in the relative gamma band power prompted us to further investigate neuronal excitability in D2J mice, which is presented in the following section.

### Cortical pyramidal neurons in D2J Scn2a^gt/gt^ mice display hyperexcitability

Given the pronounced SWDs observed in D2J *Scn2a^gt/gt^* mice via cortical EEG, we conducted whole-cell patch-clamp recordings to investigate the neuronal intrinsic firing properties underlying SWD manifestation (**Figure 3**). We focused on pyramidal neurons in the superficial layer II/III, which were located near EEG surface recording electrodes (**Figure 3A**). We noticed that neurons from the D2J *Scn2a^gt/gt^* mice fired significantly more action potentials (APs) upon a step current injection from 50–400 pA compared to the D2J WT mice (**Figure 3, B and C**). Additionally, D2J *Scn2a^gt/gt^* neurons exhibited significantly depolarized resting membrane potential (RMP) and increased input resistance, along with a decreased rheobase compared to the D2J WT neurons. Taken together, this demonstrates heightened intrinsic excitability (**Figure 3, D-I**). Notably, the APs in D2J *Scn2a^gt/gt^* mice had higher threshold potential, lower amplitude, and higher fast after-hyperpolarization (AHP), consistent with our previous results obtained in the B6 *Scn2a^gt/gt^* mice (15) (**Figure 3, J-L**). Moreover, since potassium channels can control the RMP and repolarization, the increase in AHP could indicate a major dysfunction in the K_V_ channels(51). Phase plot analysis was done by plotting the rate of MP change (i.e., 1^st^ derivative of voltage change) against the MP (mV), which could highlight different aspects of the AP more clearly (52). The representative AP traces and phase-plane plots show that the APs in D2J *Scn2a^gt/gt^* neurons exhibit lower amplitude and depolarization slope compared to the D2J WT mice. Such changes in the shape of APs suggest an overall dysfunction of Na_V_ and possibly K_V_ channels (**Figure 3, G and H**).

**Figure 3.**
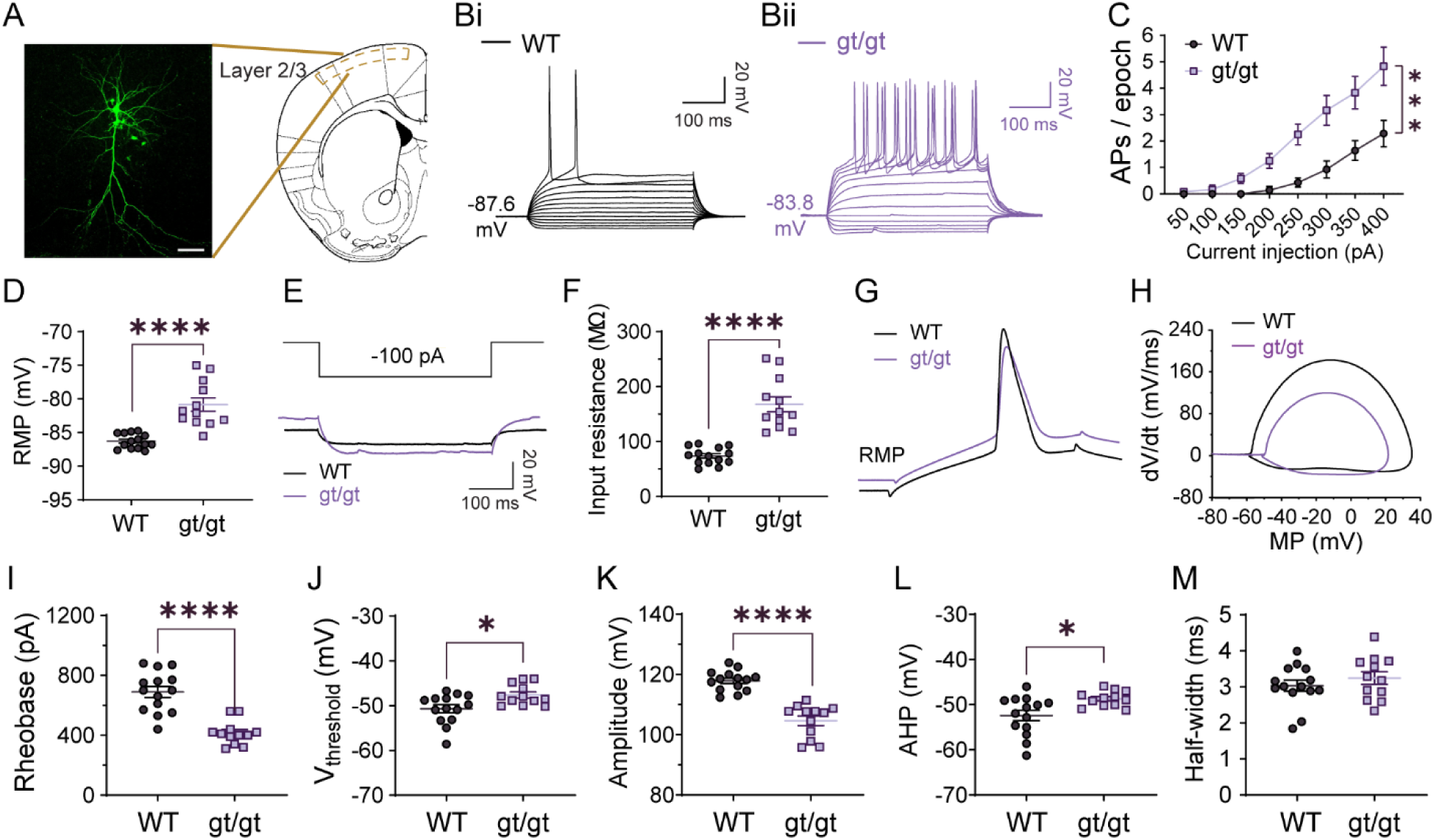
Severe *Scn2a* deficiency renders intrinsic hyperexcitability of layer 2/3 pyramidal neurons in the D2J *Scn2a^gt/gt^*mice. (A) Fluorescence imaging of a pyramidal neuron injected with biocytin after the patch clamp recording. Atlas on the right shows the location of patched cells which was in the superficial layer 2/3 of the somatosensory cortex above caudoputamen. (Bi-Bii) Representative current-clamp recordings of pyramidal neurons from D2J wild-type (WT, black) and homozygotes (*Scn2a^gt/gt^*) (violet) mice were obtained at the resting membrane potential (RMP). (C) The number of action potentials (APs) generated in response to stepwise increased current pulses was significantly higher in the D2J *Scn2a^gt/gt^* mice. (D) Pyramidal neurons in D2J *Scn2a^gt/gt^* mice had significantly higher resting membrane potential (RMP) compared to the D2J WT. (E) Representative traces in response to –100 pA injection in D2J *Scn2a^gt/gt^* and D2J WT. (F) Pyramidal neurons in D2J *Scn2a^gt/gt^* mice have significantly higher input resistance compared to D2J WT. (G) Typical AP spikes of pyramidal neurons from D2J WT (black) and *Scn2a^gt/gt^* (violet) mice were obtained at the normal RMP. (H) Example phase-plane plots show different AP shapes in D2J WT and *Scn2a^gt/gt^*. (I) The mean spike rheobase (pA) for pyramidal neurons in D2J *Scn2a^gt/gt^* mice was significantly lower than in D2J WT mice. (J) The mean voltage threshold (mV) was unchanged in D2J *Scn2a^gt/gt^* vs. WT. (K) The mean AP amplitude (mV) of pyramidal neurons in D2J *Scn2a^gt/gt^* was significantly decreased compared with D2J WT neurons. (L-M) The mean AP fast after-hyperpolarization (AHP) and half-width value of pyramidal neurons were not changed in D2J *Scn2a^gt/gt^*. Data are presented as mean ± SEM. Statistical analyses: Two-way ANOVA and unpaired two-tailed non-parametric Mann-Whitney U test for each current pulse (C). Unpaired Student’s t-test was used for all other comparisons (D)(F)(H)(I-M). *p < 0.05; **p < 0.01; ***p < 0.001; ****p < 0.0001. Exact p values can be found in Supplemental Table 1.

To determine whether the increase in RMP was the main contributor to AP firing increase, we performed the same set of recordings while holding the neuron at a fixed membrane potential of –80 mV, which was slightly more depolarized than the average RMP of – 86.62 mV in D2J WT mice (**Figure 3D**). Similar to the recordings at RMP, we still detected a significantly increased AP firing frequency in response to a step current injection, an increase in input resistance, decreased rheobase, a decrease in AP amplitude, and an increase in AP after-hyperpolarization (AHP) (**Supplemental Figure 4**). Together, these results suggest that the depolarization of RMP was not sufficient to explain the neuronal hyperexcitability.

### Global restoration of Na_V_1.2 expression in adulthood reduced short SWDs in D2J Scn2a^gt/gt^ mice

For monogenic epilepsies such as *SCN2A* disorders, correcting the defective gene is the most direct therapeutic approach. However, whether restoring *Scn2a* expression alleviates *Scn2a* deficiency-related seizures remains unknown. To address this, it is essential to assess the impact of Na_V_1.2 restoration on seizure severity in D2J *Scn2a^gt/gt^* mice. These mice carry a gene-trap cassette in their *Scn2a* flanked by two *frt* sites, which can be excised by flippase (FLP) to generate a ‘rescued’ allele (**Supplemental Figure 5A**). Given that AAV-PHP.eB can cross the blood-brain barrier and achieve whole-brain expression in various mouse strains, including DBA/2J (53), we implemented global gene-trap removal via tail vein injection of AAV-PHP.eB-Flpo. This approach has been previously validated in adult B6 *Scn2a^gt/gt^*mice, achieving partial restoration of Na_V_1.2 (15, 30).

To take advantage of the genetic construct of our transgenic mice, we recorded a 1-week video-EEG before and after tail-vein AAV-PHP.eB-Flpo injection at the adult stage (**Figure 4**). The gene-trap cassette incorporates a reporter *LacZ* gene that encodes β-galactosidase, which serves as an indicator for detecting the presence or absence of the cassette. β-galactosidase staining indicated that the Flpo incorporation successfully removed the gene-trap cassette in the whole brain, allowing restoration of *Scn2a* transcription towards the WT level (**Supplemental Figure 5C**). Additionally, the western blot analysis revealed that Na_V_1.2 expression was partially restored (∼62.15%) in D2J *Scn2a^gt/gt^*-Flpo mice (**Supplemental Figure 5B**). Notably, Flpo injection in adult D2J *Scn2a^gt/gt^* mice significantly reduced short SWD frequency: S-SWD frequency was 5.87 ± 1.22 pre-Flpo injection vs. 3.76 ± 1.02 per hour post-Flpo injection (*p < 0.05, paired t-test), though its effect on long SWDs was limited (**Figure 4, B-C**). Flpo injection also led to a significant decrease in relative theta power during the light-on period, possibly due to the considerable reduction of the S-SWD number (**Supplemental Figure 5, D-E**). Together, these results suggested that systemic restoration of Na_V_1.2 expression in adult mice could partially rescue the frequency of short spike-wave discharges in the D2J Scn2a *Scn2a^gt/gt^* mice and alter the overall EEG power.

**Figure 4.**
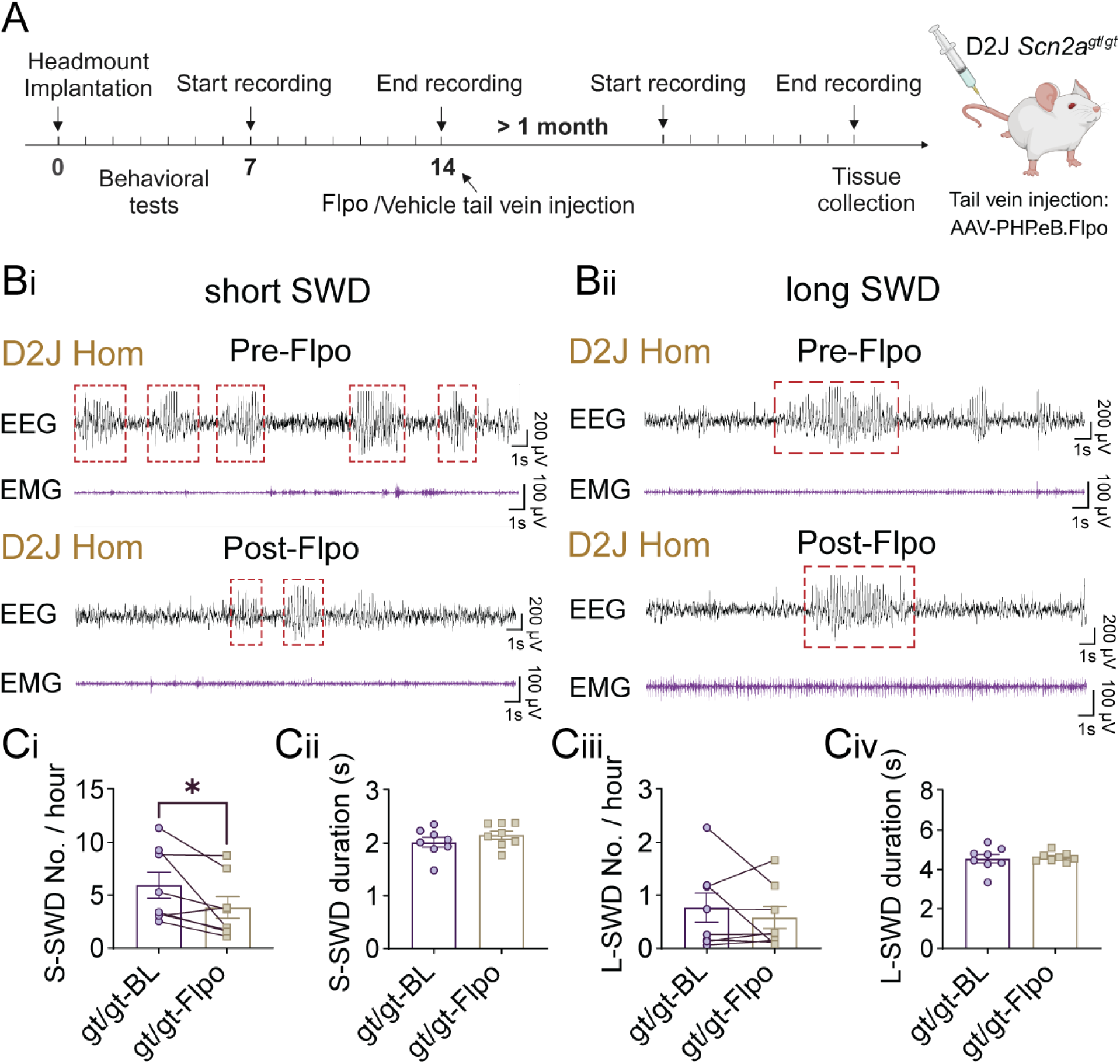
Genetic restoration of *Scn2a* expression by tail vein injection of AAV-PHP.eB-Flpo reduced short SWDs in the D2J *Scn2a^gt/gt^* mice. (A) Schematic timeline of Flpo or control virus injection. D2J *Scn2a^gt/gt^* mice underwent EEG headmount implantation and recovered in the home cage for one week. One-week continuous video-EEG recordings were conducted pre-Flpo and 1-month post-Flpo tail vein injection. (Bi-Bii) Representative EEG2 traces showing spike-wave discharges (SWDs) in the D2J *Scn2a^gt/gt^* mice before and after AAV-PHP.eB-Flpo injection. (Ci-Civ) Quantification of EEG data showed that D2J *Scn2a^gt/gt^*mice had a significant reduction in S-SWD number per hour but no change in S-SWD duration (s) or L-SWD frequency/duration after Flpo injection. Data are presented as mean ± SEM. Statistical analyses: Paired Student’s t-tests were used in all analyses (Ci-Civ)(Ei-Eiv). *p < 0.05; **p < 0.01; ***p < 0.001; ****p < 0.0001.

### RNA sequencing in D2J Scn2a^gt/gt^ mice unveils differential gene expression, highlighting downregulation of multiple potassium channels

To investigate the gene expression profile in the D2J *Scn2a^gt/gt^* mice compared to their D2J WT littermates, we performed whole cortex bulk RNA sequencing after 1-week video-EEG recording. We identified 1718 upregulated and 1984 downregulated genes and selected seizure-related gene sets from the DisGeNet library for further analysis (54). Unsupervised hierarchical clustering revealed that the overall expression profile of 616 seizure-related genes was notably different in *Scn2a^gt/gt^* mice (**Figure 5A**). Many of the downregulated epilepsy-associated genes encode functional proteins that regulate neuronal excitability, whose disturbance contributes to pathological synchronization leading to seizures. As expected, the *Scn2a* gene was most significantly downregulated in the D2J *Scn2a^gt/gt^* mice compared to the D2J WT mice in the volcano plot (**Figure 5B, red dashed box**). Notably, *Scn8a* (encoding the Na_V_1.6 channel) and *Scn1b* (encoding the Na_V_β-1 subunit) were significantly downregulated (**Figure 5B**). When comparing the bulk RNA-seq data for D2J mice to the B6 mice reported in the previous paper (29), we noticed a highly similar global downregulation of potassium channels, including K_V_. This result suggests that compensatory potassium channel downregulation is a robust and conserved phenotype in *Scn2a^gt/gt^* mice regardless of their strain background (**Figure 5C**). In contrast, there are strain-specific differential gene alterations. For instance, a group of synaptic-related genes was significantly decreased in the D2J *Scn2a^gt/gt^* mice but not affected in the B6 mice, suggesting that specific synaptic genes were uniquely altered in response to *Scn2a* deficiency depending on the strain (**Figure 5D**). Likewise, multiple calcium channel-related genes were significantly down-regulated in the D2J *Scn2a^gt/gt^* mice while essentially unchanged in the B6 mice (**Figure 5E**). In particular, this includes *Cacng2* and *Cacna1a*, genes that play critical roles in the pathogenesis of absence epilepsy (55, 56). There are also several glutamate-, GABA-, myelin-, and growth-related genes that were significantly downregulated in D2J but not B6 mice, again emphasizing the contribution of strain difference to the EEG and behavioral phenotypes in mice (**Supplemental Figure 6, A-D**).

**Figure 5.**
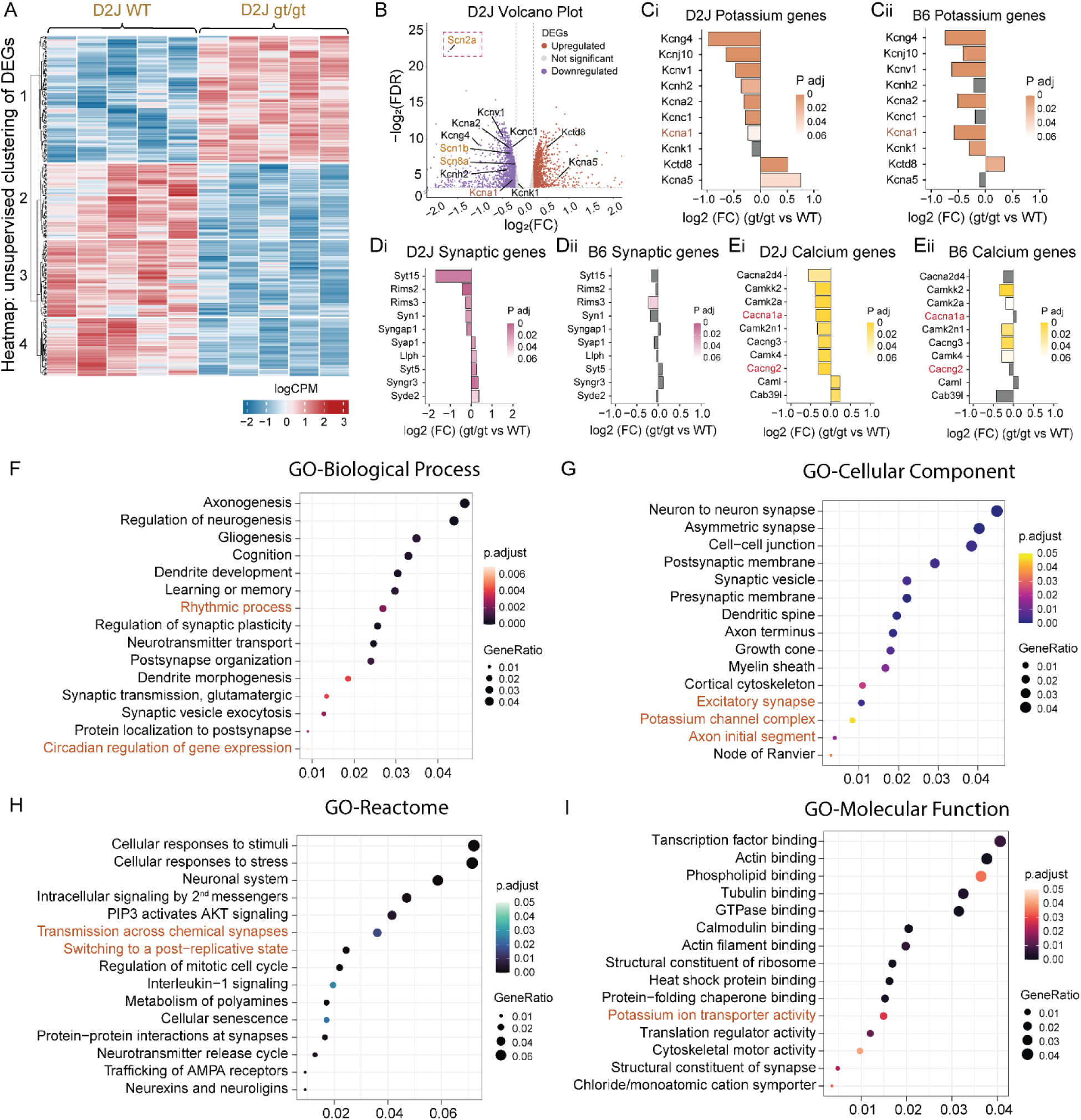
Bulk RNA-seq analyses revealed possible molecular mechanisms underlying the seizure susceptibility in D2J *Scn2a^gt/gt^* mice including reduced expression of K_V_ channels. (A) Heatmap showing an overall pattern of unsupervised clustering of seizure-related differentially expressed genes (DEGs) in the D2J WT vs. D2J *Scn2a^gt/gt^*. logCPM: log counts per million. (B) Volcano plot showing the –log_2_(FDR) and log_2_(FC) of voltage-gated sodium and potassium channels in D2J WT vs. D2J *Scn2a^gt/gt^* cortices. Note that the *Scn2a* gene was significantly downregulated as expected (red dashed box). (C-E) Side-by-side comparison of potassium channel genes, synaptic associated genes, and calcium channel genes that were significantly downregulated in the D2J *Scn2a^gt/gt^* mice with the same gene changes in the B6 based on previous study. (F-I) Gene ontology analyses based on the biological process, cellular component, Reactome, and molecular function of the D2J *Scn2a^gt/gt^* mice compared to D2J WT mice. FDR: False discovery rate; FC: Fold change. Data are presented as mean ± SEM. Statistical analyses: Please refer to the methods section for specific statistical analysis used for the bulk RNA seq data. Multiple hypothesis testing correction was done using the Benjamini-Hochberg method, with genes showing an FDR below 0.1 considered differentially expressed. Exact p values can be found in Supplemental Table 1.

Gene ontology (GO) analyses revealed global functional changes in the D2J *Scn2a^gt/gt^* mice (**Figure 5, F-I**). As expected, neurodevelopment-related pathways such as synaptic transmission, axogenesis, neurogenesis, and gliogenesis were most substantially altered (**Figure 5, F-G**). Additionally, genes that regulate circadian rhythm were affected in the D2J *Scn2a^gt/gt^* mice, a result consistent with our previous study on B6 *Scn2a^gt/gt^* mice (29) and in line with the EEG power spectral pattern (**Supplemental Figure 2**). Note that potassium channel activity was again significantly altered in the GO-cellular component and GO-molecular function analysis (**Figure 5, G and I**). Overall, this set of GO data demonstrated that synaptic and neurodevelopment-related functions were most significantly altered in the D2J *Scn2a^gt/gt^* mice.

### Delivery of exogenous human K_V_1.1 via AAV rescued both short and long SWDs in the D2J Scn2a^gt/gt^ mice

Alterations in neuronal action potentials and RNA sequencing results suggest that the downregulation of voltage-gated potassium channels (K_V_) expressions may contribute to the pathophysiology of absence seizures in *Scn2a*-deficient mice (**Figures 3 and 5**). Accordingly, we investigated whether introducing K_V_ channels could reduce epileptiform activity in D2J *Scn2a^gt/gt^* mice. Exogenous expression of human K_V_1.1 (encoded by *Kcna1*) has recently been proposed as a novel gene therapy strategy for treating refractory epilepsies (27, 28, 57). Since we noticed prominent downregulation of potassium channels in both D2J and B6 *Scn2a^gt/gt^* mice (**Figure 5, B and C**) (15), we conducted experiments to test if this potential targeted gene therapy approach could reduce the prominent absence seizure observed in D2J *Scn2a^gt/gt^* mice (**Figure 6**). We bilaterally injected an AAV9-hK_V_1.1 vector (**Supplemental Table 1**) into the lateral ventricles of the D2J WT and *Scn2a^gt/gt^* mice (**Figure 6A**). The virus was constructed with a *Camk2a* promoter-driven Tet-On system and thus can be activated by a doxycycline diet (**Figure 6B**). We then recorded the EEG signal of the same mouse before and after 1-month doxycycline induction (**Figure 6A**). We discovered that exogenous expression of human K_V_1.1 protein during the adult stage in *Scn2a^gt/gt^* mice was able to alleviate the number of both short and long SWDs: the S-SWD in D2J *Scn2a^gt/gt^*mice was 5.84 ± 1.68 per hour during baseline which reduced to 2.55 ± 1.24 per hour post-doxycycline induction, [BL-Dox] *Scn2a^gt/gt^* adjusted ***p < 0.001, two-way ANOVA (**Figure 6Cii)**; Similarly, the L-SWD events with a frequency of 0.46 ± 0.16 per hour was reduced to 0.24 ± 0.10 per hour post-doxycycline, [BL-Dox] *Scn2a^gt/gt^* adjusted *p < 0.05 (**Figure 6Dii)**. The duration of SWDs in D2J *Scn2a^gt/gt^* mice remained largely unchanged (**Figure 6, Ciii and Diii**). The D2J WT mice were not significantly affected by hK_V_1.1 overexpression. These mice appear active and alert with low SWD frequency (**Figure 6, Cii-Ciii and Dii-Diii**).

**Figure 6.**
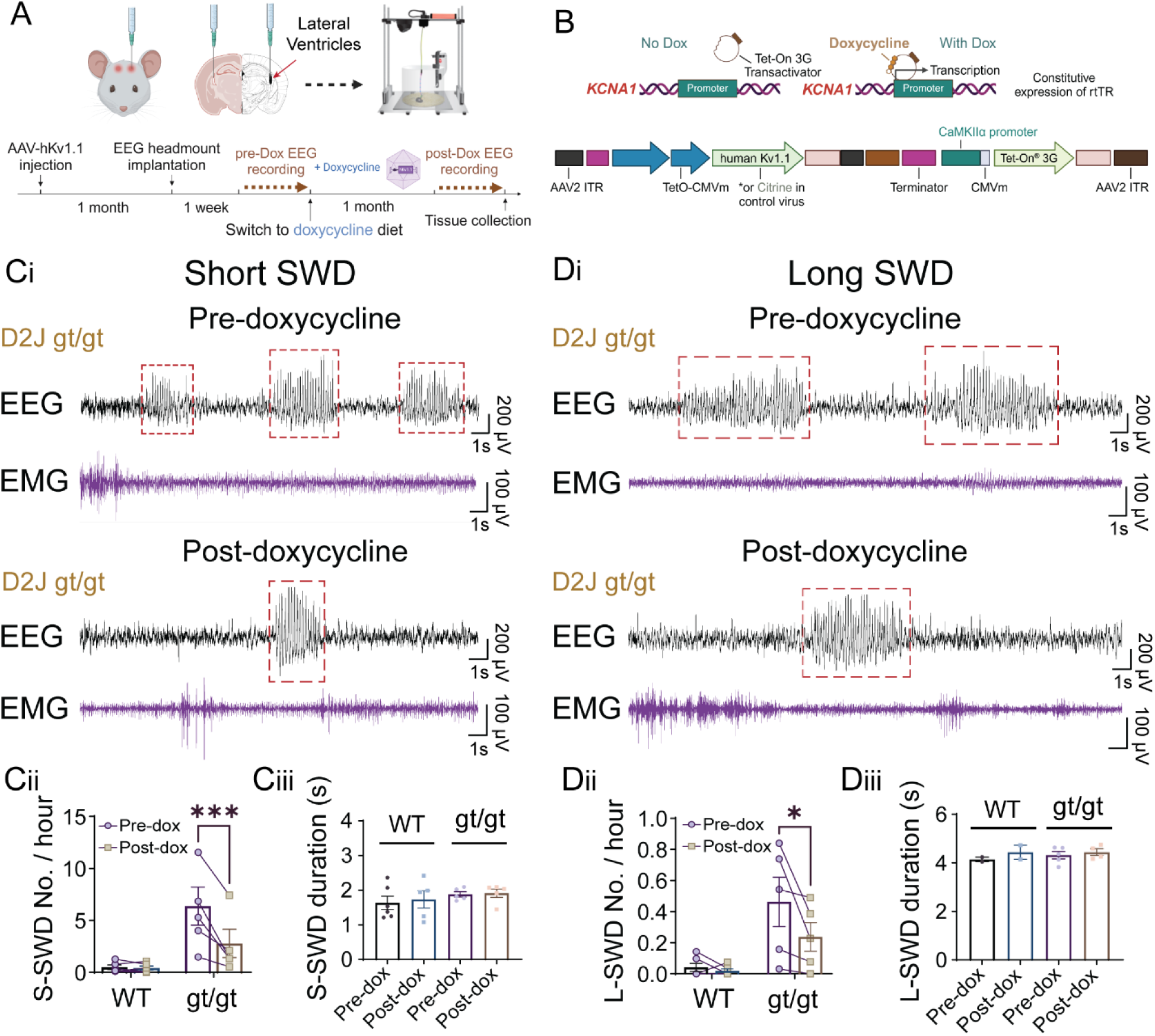
Overexpression of human K_V_1.1 in the D2J *Scn2a^gt/gt^* mice reduced the frequency of both short and long SWDs related to absence seizures. (A) Schematic of the AAV-hK_V_1.1 virus injection experiment and video-EEG recording timeline. (B) Construct of the AAV-hK_V_1.1 virus showing the addition of doxycycline activating the genetic expression under a Tet-On system. (Ci-iii) Example EEG traces and quantifications showing a significant reduction in SWD frequency for D2J *Scn2a^gt/gt^* after doxycycline induction of the AAV-hK_V_1.1 virus expression. The duration of SWDs was unchanged. (Di-iii) Example EEG traces and quantifications showing a significant reduction in absence seizures frequency for D2J *Scn2a^gt/gt^*after doxycycline induction of the AAV-hK_V_1.1 virus expression. The duration of absence seizures was unchanged. Data are presented as mean ± SEM. Statistical analysis: Paired Student’s t-tests were used in all analyses (Cii-Ciii)(Dii-Diii). *p < 0.05; **p < 0.01; ***p < 0.001; ****p < 0.0001. Exact p values can be found in Supplemental Table 1.

To assess the translational potential of AAV-hK_V_1.1, we generated human cerebral organoids with *SCN2A* deficiency. Consistent with the aforementioned findings, we observed substantial downregulation of multiple K_V_ channel genes, particularly *KCNA1*, in human neurons. AAV-hK_V_1.1 transduction effectively elevated *KCNA1* expression and reduced neuronal excitability in human brain organoids (**Supplemental Figure 7**). Collectively, these findings suggest that exogenous hK_V_1.1 expression may represent a potential targeted gene therapy for seizures associated with *SCN2A* deficiency.

## Discussion

In this study, we discovered that adult *Scn2a*-deficient gene-trap (*Scn2a^gt/gt^*) mice in the ‘seizure-prone’ DBA/2J (D2J) background exhibited more prominent absence seizures than those in the ‘seizure-resistant’ C57BL/6N (B6) background. Using congenic D2J *Scn2a^gt/gt^* mice as a preclinical disease model, we found that adult restoration of Na_V_1.2 was able to decrease spike-wave discharge (SWD) numbers. RNA sequencing reveals that *Scn2a^gt/gt^* mice show differential gene expression, which indicates potential mechanisms behind strain-dependent seizure susceptibility difference. Importantly, significant downregulation of voltage-gated potassium channels (K_V_) was observed in both strains of *Scn2a^gt/gt^*mice, indicating a conserved compensatory pathway. Employing K_V_ as an alternative target, we discovered that expression of the exogenous human K_V_1.1 protein significantly rescued the absence seizure phenotype, demonstrating the potential of gene therapy in treating *Scn2a* deficiency-related epilepsy.

In C57BL/6 mice with haplodeficient Na_V_1.2, prior studies have suggested a relatively mild increased number of SWDs compared to the WT (12, 58). Evaluating therapeutic interventions requires a robust phenotype with a broad dynamic range to accurately assess efficacy, suggesting the need for an enhanced disease model. Since mouse genomic background has a substantial effect on seizure susceptibility, we enhanced the seizure phenotype by rederiving *Scn2a^gt/gt^* mice in the ‘seizure-susceptible’ D2J strain (**Figure 1**) (18, 23). Despite prominent absence seizures and abnormal behaviors, no spontaneous tonic-clonic seizures were observed in adult D2J *Scn2a^gt/gt^* mice. Similar seizure phenotypes have been observed in other *Scn* mouse models. For example, *Scn8a*-null mice exhibited only spontaneous absence seizures (21), and *Scn3a*-null mice showed no spontaneous convulsive seizures (59). In contrast, *Scn1a* (*60*) and *Scn1b* (*61*) haplodeficient mice display unprovoked tonic-clonic seizures at juvenile age. Distinct seizure phenotypes associated with Na_V_ isoform deficiencies in mice may reflect neuronal subtype-specific expression: Na_V_1.2, Na_V_1.3, and Na_V_1.6 primarily affect excitatory neurons, while Na_V_1.1 and Na_V_β1 impact inhibitory neurons (62, 63). However, such hypotheses warrant further investigation, and ongoing studies are examining the roles of Na_V_ isoforms in interneurons (64). Phenotypic differences between humans and rodents may also arise from disparities in neural network scale and Na_V_ channel function patterns across neuronal subtypes during development. Future studies are needed to elucidate the mechanism behind epileptogenesis in these models and their implications in human sodium channel*-*related diseases.

Patients with *SCN2A* mutations exhibit considerable clinical heterogeneity. This variation is partly due to differences in individual genomic profiles, with certain genetic predispositions leading to more severe seizures. Likewise, different mouse strains carry distinct genetic backgrounds, and polymorphisms in specific genes contribute to seizure susceptibility, partially mirroring the genomic variations observed in humans. For instance, fine-mapping has identified candidate modifier genes underlying strain-dependent epilepsy differences in a *Scn1a* mouse model of Dravet syndrome (65). To investigate strain-dependent seizure severity, we compared RNA-seq data from D2J mice with published data for B6 mice to assess differential gene expression patterns (29). We discovered multiple epilepsy-related genes that are significantly downregulated in D2J mice but remain unchanged in B6 mice (**Figure 5 and Supplemental Figure 6**). For example, *Gabra2*, which encodes a GABA_A_ receptor subunit, functions as a genetic modifier in *Scn1a*- and *Scn8a*-associated developmental and epileptic encephalopathy (DEE) and is linked to strain-dependent seizure susceptibility (66, 67); voltage-gated calcium channel genes such as *Cacna1a* and *Cacng2* are strongly associated with absence epilepsy (55, 56) and has close interaction with many sodium channel genes (68, 69). These findings indicate that, compared to B6 mice, D2J mice exhibit distinct global gene expression alterations in response to germline Na_V_1.2 deficiency, leading to exacerbated epileptiform discharges. Nevertheless, our experiment did not rule out important contributions from single nucleotide polymorphisms (SNPs) in DJ2 and B6 mice, as investigated in other strain difference studies using quantitative trait locus (QTL) fine-mapping (70). Examining differential gene contribution to epileptogenesis in these strains could potentially provide additional insights into the heterogeneity observed in human patients, fostering the development of targeted precision medicine.

Since we recorded EEG for mice before bulk RNA sequencing, it allows us to correlate the absence seizure severity (ranked based on overall SWD frequency/duration) with the normalized counts for genes of interest. We first plotted the *Scn2a* count against the *Kcna1* count, which showed no significant correlation (**Supplemental Figure 6E**). However, we found a significant negative correlation between *Scn2a* count and seizure severity, as well as *Kcna1* count and seizure severity (**Supplemental Figure 6, F and G**). This suggests that deficiencies in either of these genes likely contribute to the absence seizure phenotype. In contrast, not every significantly downregulated gene is correlated with absence seizure severity. For instance, although we noticed a significant downregulation of *Cacng2* in the bulk-RNA seq, its expression was not significantly correlated with absence seizure severity, indicating that it might be a ‘risk factor’ predisposing the D2J mice to have SWDs but not a key factor contributing to the seizure severity in *Scn2a* deficiency (**Supplemental Figure 6H**).

Recently, many gene therapies have been tested *in vivo* to advance the treatment of monogenic epilepsies (25). For instance, antisense oligonucleotide (ASO)- and viral vector-mediated channel restoration have been demonstrated to reduce seizure pathology in multiple mouse models of DEE (71–73). Although achieving physiological Na_V_1.2 expression in *SCN2A* LoF patients still presents significant technical challenges, our gene-trap transgenic mice offer a proof-of-concept platform. This model enables the evaluation of seizure outcomes following the global restoration of *Scn2a* expression by removing the trapping cassette. Despite relatively low plasticity in the adult brain, EEG recording showed that tail vein injection of AAV-PHP.eB-Flpo significantly reduced the number of short SWDs in D2J *Scn2a^gt/gt^*mice (**Figure 4**). This encouraging finding demonstrates the potential of reducing *Scn2a* LoF-related seizures at a later stage through systemic AAV-mediated upregulation of *Scn2a* expression. Nevertheless, it is important to acknowledge that selection of the treatment window is crucial in developmental epilepsies, as recent mouse studies on *SCN1A* and *SCN1B* Dravet syndrome discovered that only neonatal gene therapy effectively alleviates sudden unexpected death in epilepsy (SUDEP) from convulsive seizures (74, 75). Although Flpo injection in adulthood reduced S-SWDs, the effect of Na_V_1.2 restoration is limited since neither long SWDs (>3.5 s) nor SWD duration was significantly reduced (**Figure 4**). Hence, it is possible that earlier intervention could further suppress absence seizures.

In our RNA sequencing analysis, we identified a set of potassium channel genes consistently downregulated in *Scn2a^gt/gt^* mice across both D2J and B6 strains. This observation was further confirmed in *SCN2A*-deficient human cerebral organoids, indicating a compensatory K_V_ channel reduction in response to severe Na_V_1.2 deficiency, which is likely conserved across mouse strains and species (**Figure 5C, Supplemental Figure 6D, and Supplemental Figure 7**). We therefore explored K_V_ as a potential therapeutic target in addition to directly restoring Na_V_1.2 expression (28, 76). Notably, *KCNA1* has been proposed as a promising therapeutic target for refractory epilepsies and has demonstrated efficacy in mouse models of visual cortex epilepsy (27), temporal lobe epilepsy (27), focal cortical dysplasia (57), and focal neocortical epilepsy (77). By incorporating a tetracycline-dependent gene transcriptional design, we were able to express exogenous human K_V_1.1 through doxycycline induction (78). It is worth noting that the introduction of K_V_ to inhibitory interneurons may reduce their excitability, leading to neural network disinhibition, which could potentially worsen seizures in *Scn2a^gt/gt^*mice. Therefore, the *KCNA1* gene expression was designed with an excitatory neuron-specific *CaMKII*α promoter, which reduces unwanted expression in the GABAergic inhibitory neurons. This AAV-mediated human K_V_1.1 transgene was delivered in the adult mouse brain through intracerebroventricular (ICV) injection (**Figure 6, A-B**). Even at the adult stage, this approach successfully reduced both short and long SWDs in the D2J *Scn2a^gt/gt^* mice (**Figure 6**). Additionally, application of the same AAV-hK_V_1.1 significantly elevated *KCNA1* expression and effectively reduced neuronal firing in human brain organoids, reinforcing the potential of K_V_1.1 as a therapeutic target for *SCN2A* deficiency-related epilepsies (**Supplemental Figure 7**).

In conclusion, our study established a unique *Scn2a* deficiency-related epilepsies disease model for testing new therapeutics. This animal model allows us to assess the treatment efficacy and route of delivery of gene therapies, offering valuable insight into future clinical translation. Additionally, we examined gene expression patterns that may underlie the strain-dependent differences in absence seizure severity between B6 and D2J *Scn2a^gt/gt^* mice. These findings aim to expand our understanding of the *SCN2A* disease mechanism and help pave the way for genetic interventions to treat epilepsy in patients with *SCN2A* LoF mutations.

## Methods

### Experimental animals

All experimental procedures were approved by the Purdue University Institutional Animal Care and Use Committee (IACUC) and conducted according to ethical guidelines provided by the NIH and AAALAC International. All mice were bred in the Purdue animal facility and both sexes were used in equal proportion. Mice were housed in a maximum of five per cage under a 12:12 h light/dark cycle with *ad libitum* access to food and water. The animal room was maintained at a consistent temperature (68°F to 79°F) and humidity (30% to 70%) based on the USDA Animal Welfare Regulations (AWR) and the ILAR Guide for the Care and Use of Laboratory Animals. For all surgeries, mice were administered analgesic buprenorphine based on Purdue Animal Care Guideline to assist with recovery.

C57BL/6N-*Scn2a1^tm1aNarl^*/Narl mice generated previously by the lab were used in this study(14). The *Scn2a* gene in the *Scn2a^gt/gt^* mice contains a gene-trap cassette that includes two *frt* sites, strong splicing acceptors, and a reporter gene *LacZ* (encoding the β-galactosidase enzyme). To produce the *Scn2a^gt/gt^* congenic mice in the DBA/2J strain background, C57BL/6N- *Scn2a^WT/gt^* (B6-Het) mice were backcrossed to the inbred DBA/2J WT mice purchased from Jackson Laboratory^®^ (RRID:IMSR_JAX:000671) for 8 generations. The genomes of the resulting congenic mice were validated through Giga Mouse Universal Genotyping Array (GigaMUGA) with >99% identity compared to the DBA/2J WT inbred mice from JAX. Then, DBA/2J-*Scn2a^WT/gt^*mice were crossed (D2J-Het x D2J-Het) to create an in-house colony for experiments in this study (i.e. D2J-*Scn2a^gt/gt^* and D2J-WT).

### Genotyping

At weaning (21-28 days old), mice from the colony were identified via ear punch, and the ear tissues were collected for genotyping. DNA was extracted by heating the tissues in 50 mM NaOH followed by the addition of 1M Tris (pH = 8) and centrifugation of 12,000 g for 10 min. The desired DNA segment was amplified using gene-specific polymerase chain reaction (PCR) with primers (see materials table) and segregated via agarose gel electrophoresis. The PCR product of the wild-type allele is 240 base pairs (bp) and the tm1a (gt) allele’s PCR product is 340 bp. The heterozygotes show two bands at 240 bp and 340 bp.

### EEG surgeries and recordings

All procedures were conducted according to the EEG surgical guide provided by Pinnacle Technology. Adult mice were anesthetized by intraperitoneal injection of a mixture of ketamine/xylazine (100/10 mg/kg body weight) dissolved in sterile saline. The scalp surface was exposed, and the prefabricated 2EEG/1EMG mouse headmount (Cat: 8201) was implanted on the skull with the front edge placed 3–3.5 mm anterior of the bregma. Pilot holes were drilled with a 23G needle and four 0.1’’ stainless steel screw electrodes were inserted in the cortex. The contact between the recording electrodes and the headmount was secured by silver epoxy and then covered with dental acrylic. The two EMG probes were embedded into the nuchal muscles. The continuity between each bipolar electrode and its corresponding metal contacts was tested by a multimeter. After surgery, the animals were returned to their home cages to recover for at least one week.

Continuous synchronized video-EEG/EMG were recorded 24/7 for one week using the Pinnacle Sirenia^®^ Acquisition System. The signals were captured using a pre-amplifier with a gain of 100 Hz (Catalog: 8202) connected to a data conditioning and acquisition system (8206-HR) through a 3-channel mouse commutator/swivel (6-Pin) (8204–723) at a 400 Hz sampling frequency with a 100 Hz lowpass filter. EEGs were time-synchronized with continuous video recordings from IP cameras with automated IR sources.

### Epileptiform discharges and power spectral analysis

Each seizure event was first screened by the Sirenia SeizurePro^®^ software based on power and then manually verified by trained blinded observers in combination with the corresponding video recordings. Spike-wave discharges (SWDs) were identified using criteria established for analyzing mouse models of absence epilepsy. In brief, SWDs were defined as rhythmic biphasic synchronous spike-and-wave complexes (5–7 Hz) with a duration of >1 s and discharge amplitude at least twofold higher than the average nearby baseline voltage with concomitant video-recorded behavioral arrest (33, 79). We noticed that in our EEG cortical recordings using screw electrodes wild-type D2J mice typically do not have SWDs longer than 3.5 s. Therefore, to better characterize the EEG phenotype in these animals, we divided the SWDs into short (S-SWD; 1–3.5s) and long (L-SWD; >3.5 s) episodes, where the long SWDs more closely resemble absence seizures and are defined by sudden behavioral arrest, fixed staring posture, and bilateral synchronous SWDs lasting more than 3.5 s (31).

Video recordings and the EMG trace during the SWD events were checked to confirm the sudden behavioral arrest or loss of consciousness associated with SWD and to exclude artifacts from muscle activity such as drinking water and grooming (80). The ‘SWD clusters’ were defined as five or more SWD episodes that occurred with an inter-episode interval of maximal 60 s (81). Myoclonic seizures were identified first based on a significant amplitude increase in EEG1 and EMG traces, and then videos were inspected to identify sudden jumps, wild running, and myoclonic jerks (81). Since the anterior EEG2 signal was much stronger than the posterior EEG1 signal, we used the EEG2 signal as the readout of absence seizures for the rest of the study.

Power spectra were calculated for light-on and light-off periods separately by Fast Fourier Transform (FFT) with Hann (cosine-bell) data window set using an epoch of 10s based on the following frequency bands: full (0–100 Hz), delta (0.5–4 Hz), theta (4–8 Hz), alpha (8–13 Hz), beta (13–30 Hz), and gamma (30–100 Hz). The relative power was calculated by dividing each power band with the full power (0–100 Hz).

### Adeno-Associated Virus (AAV) Production

pAAV-EF1a-mCherry-IRES-Flpo was a gift from Karl Deisseroth (Fenno et al., 2014) (Addgene plasmid # 55634; http://n2t.net/addgene:55634; RRID: Addgene_55634), AAV9-PHP.eB-EF1a-mCherry-IRES-Flpo with the titer of 2.56×10^13^ GC/mL was packaged by the Penn Vector Core; Control virus, PHP.eB-Ef1a-DO-mCherry-WPRE-pA with the titer of 1.2×10^13^ GC/mL was packaged by Bio-Detail Corporation. K_V_1.1-Negative control: AAV9/NegCTRLCam with the titer of 2.75×10^13^ GC/mL; K_V_1.1-Negative fluorescence control: AAV9/TOCitPCamTA with the titer of 1.70×10^13^ GC/mL; and K_V_1.1-Positive: AAV9/TOK_V_1CamTA with the titer of 1.50×10^13^ GC/mL were packaged by the Horae Gene Therapy Center.

### Systemic and stereotaxic AAV injection and doxycycline activation

The *Scn2a^gt/gt^* mice contain a gene-trap cassette flanked by two *frt* sites. The *frt* sites can be recognized by the flippase (FLP), leading to the removal of the trapping cassette, essentially resulting in a ‘rescue allele’. Tail-vein injection of the blood-brain-barrier-crossing AAV-PHP.eB-Flpo vector is expected to globally remove the gene-trap cassette and yield a “rescue allele” with a full-length *Scn2a* transcript (14, 82). To globally restore *Scn2a* transcription, each adult mouse received 5 × 10^11^ genome copies (GC) of Flpo or control AAV via tail vein injection to achieve systemic delivery.

For the viral injection into lateral ventricles through cerebral spinal fluid circulation, mice were anesthetized with ketamine/xylazine (100/10 mg/kg, i.p.) and secured in a stereotaxic apparatus with ear bars (RWD Ltd, China). After exposing the skull via a small incision, small holes for each hemisphere were drilled for injection based on coordinates to bregma. Mice were bilaterally injected with AAV9/TOK_V_1CamTA or AAV9/TOCitPCamTA virus (5 × 10^12^ GC/mL with PBS) into the lateral ventricles (coordinates of the injection sites relative to bregma: AP -0.50 mm, ML ± 1.00 mm, DV -2.00 mm, 10 μL per site, at the speed of 1 μL/min) with sharpened glass pipettes (Sutter Instrument), self-made to have a bevel of 35° and an opening of 20-mm diameter at the tip (83), attached to syringe needles (200-mm diameter). The pipette was filled from the back end with mineral oil and attached to a syringe needle mounted in a microinjection syringe pump (World Precision Instruments, UMP3T-2). Before injection, the viral suspension was suctioned through the tip of the pipette. The skull over the target coordinates was thinned with a drill and punctured with the tip of the pipette. The pipette was inserted slowly (120 mm/min) to the desired depth. The virus was slowly (∼100–150 nL/min) injected into the desired location. Before being retracted out of the brain, the pipette was left in the same place for 10 min when the injection was finished. The accurate location of injection sites and viral infectivity were confirmed in mice *post hoc* by imaging sections containing the relevant brain regions.

Animals were allowed to recover from surgery for at least one week and their health condition was closely monitored during recovery. Mice were fed with control diet in the meantime. After recovery, a one-week baseline video-EEG recording was performed. Then, the mouse diet was switched to a chow that contained 200 mg/kg of doxycycline. The virus was allowed to be activated for one month followed by post-doxycycline one-week video-EEG recording to compare brain activity before and after hK_V_1.1 overexpression.

### Open field test

Mice were habituated to the scent of the researcher for 5 days before the test date. On the date of the test, mice were transferred to the behavior room 20 minutes prior to the time of the test. Mice were then placed in an open field box with dimensions 40 × 40 × 40 cm (Maze Engineers, Boston, MA) for 10 minutes at 60 lux. The center was defined as a 20 × 20 cm square in the middle of the field. Distance traveled, center duration, and velocity were recorded by EthoVision XT (Noldus, Leesburg, VA).

### Lac-Z (β-galactosidase) histology staining

For LacZ (β-galactosidase) staining, mice were transcardially perfused with cold PBS, then 2% PFA + 0.2% glutaraldehyde. The whole brains were extracted and post-fixed in 2% PFA + 0.2% glutaraldehyde overnight, followed by 72-hour 15% and then 30% sucrose dehydration. Brains were embedded in Tissue-Tek^®^ O.C.T, frozen in 2-methylbutane in dry ice, and stored in a – 80°C freezer. 25 μm thick sagittal slices were cryosectioned, and washed for 5 min in PBS followed by 10 min in PBS with 0.02% Triton X-100. The free-floating tissues were then incubated with 500 μL of freshly prepared staining solution [X-Gal solution added into Iron Buffer (1/19, v/v) and mixed thoroughly for 10 min, and incubated for 30 min at 37°C until tissues were stained blue. Specimens were washed thrice with PBS, mounted in 70% glycerol, and sealed with nail polish before storage in a 4°C fridge. Images were captured under a light microscope and analyzed using the Fiji software.

### Immunofluorescence staining

hiPSC-derived brain organoids were fixed in 4% PFA, transferred to 30% sucrose-PBS for 3 days, and embedded in a 1:1 mixture of optimal cutting temperature (OCT) compound and 30% sucrose-PBS. Cryosections (40 μm thickness) were permeabilized and blocked in 0.5% Triton X-100 and 5% normal goat serum in PBS for 1 hour at room temperature (RT). Sections were treated with primary antibodies overnight at 4°C, washed 3 × 10 mins with PBS, and then incubated with fluorophore-conjugated secondary antibodies for 1 hour at RT. After PBS wash, sections were mounted with DAPI-containing Antifade Mounting Medium and sealed with glass coverslips. Images were acquired using an LSM900 confocal fluorescence microscope equipped with an air scan module.

### Bulk-RNA sequencing RNA extraction

Six (3M+3F) *Scn2a^gtKO/gtKO^* (GT/GT) and six (3M+3F) *Scn2a^+/+^* littermate mice in the DBA/2J background were used to extract total RNA. Mice were anesthetized and transcardially perfused with ice-cold RNAse-free PBS (Boston BioProducts). The brain was removed from the skull, and the cortices were rapidly dissected, snap-frozen in liquid nitrogen, and stored at – 80°C until use. To stabilize RNA, 100 mg (tissue weight)/ml RNAlater^®^-ICE was added to the tubes, and tissues were allowed to thaw at –20°C for at least one day. Polyadenylated (Poly(A)+) RNA was isolated from 100–250 ng of total RNA using the NEBNext® Poly(A) mRNA Magnetic Isolation Module (New England Biolabs). For 5-month-old organoids, RNA was extracted from three WT lines, three heterozygous lines, and three homozygous lines. RNA extraction was performed using the QIAGEN RNeasy^®^ Mini kit according to the manufacturer’s instructions. RNA quality was checked using an Agilent TapeStation RNA ScreenTape to ensure all samples had RNA integrity numbers greater than 8. Poly-A RNA was isolated using a NEBNext Poly(A) mRNA Magnetic Isolation Module and libraries constructed using an xGen RNA Library Prep Kit (Integrated DNA Technologies) with xGen UDI Primer Pairs. Libraries were pooled and sequenced on an Element Biosciences AVITI^TM^ System using a CloudBreak FreeStyle 2 x 150 Kit (Medium Output) with 500 million reads.

### RNA-seq data analysis

Datasets were processed using fastp (v0.23.2) to remove adapter sequences and trim low-quality bases below Q30 (84). Reads longer than 50 bp after trimming were retained for further analysis. The trimmed reads were then aligned to the Mus musculus DBA_2J_v1 (85) reference genome from Ensembl release 112 (86) using the STAR Aligner (v2.7.11b)(87) in two-pass mode. Read assignment to genomic features was performed using featureCounts (v2.0.1) (88) in paired-end and reverse-stranded mode. Samples with fewer than 20 million reads mapped to features were excluded (WT_8), and an additional sample (GT/GT_6) was removed following exploratory analysis using DESeq2 (v1.34.0) in R (v4.1.3) (89).

Sex-specific stratification was observed among the samples. To account for potential sources of variation, RUVSeq (v1.28.0) (90) was used to estimate variation factors. The raw counts matrix was filtered to retain genes with at least 5 reads in two samples and normalized with the upper-quartile method via the betweenLaneNormalization() function. Generalized linear model (GLM) regression on the counts, using GT/GT vs. WT as covariates at k = 3, incorporated RUVSeq factors into the design matrix for differential expression analysis using edgeR’s (v3.36.0) (91) quasi-likelihood negative binomial model. Multiple hypothesis testing correction was done using the Benjamini-Hochberg method, with genes showing an FDR below 0.1 considered differentially expressed. These genes were analyzed for over-represented KEGG and Reactome pathways, as well as Gene Ontology terms, using clusterProfiler (v4.10.0) in R (v4.3.2) (92), with all detected genes as the background following RUVSeq analyses. Enriched pathways and Gene Ontologies were visualized using dot plots and network plots.

Log-transformed counts per million (logCPM) values were extracted with edgeR. The DisGeNet gene set library was retrieved from Enrichr (August 18, 2024) (54), and terms related to ‘Epilepsy’ were selected. Gene clustering was performed with the ComplexHeatmap package (v2.14.0) in R (v4.2.1) (93), using k-means partitioning to group genes into four clusters based on their expression patterns.

### Western blot

Mice were anesthetized and perfused with ice-cold PBS. Whole brain tissues were collected and snap-frozen to be kept at –80°C until use. Brain tissues were homogenized in 100 mg tissue/mL ice-cold N-PER™ Neuronal Protein Extraction Reagent (Thermo Fisher Scientific, 87792) or RIPA buffer (Thermo Fisher, 89901) supplemented with 1:100 protease and phosphatase inhibitors (Thermo Fisher Scientific, A32953), sonicated on ice, and centrifuged (10,000 × g, 20 min, at 4°C). The resulting supernatants were collected, and protein concentration was determined by Pierce™ BCA Protein Assay Kits. Protein volumes were adjusted based on the concentrations and boiled in Laemmli SDS-Sample Buffer (Boston BioProducts #BP-110R) at 95°C for 5 min. For electrophoresis, 40 mg of total proteins were loaded onto the 5%-8% sodium dodecyl sulfate-polyacrylamide (SDS-PAGE) gels in Tris/Glycine/SDS Electrophoresis Buffer (#1610772) and transferred onto PVDF membrane (pore size 0.45 μm) in cold Tris/Glycine Buffer (#1610771). The resulting blots were blocked in 5% nonfat milk in Tris-buffered saline with 0.1% Tween 20 (TBST) for 1 h at room temperature and incubated with the primary antibody (1:500 Rb-Na_V_1.2, Alomone ASC-002; 1:1000 Ms-bActin Invitrogen BA3R) in LI-COR Intercept Antibody Diluent with gentle nutation overnight at 4°C. The next day, the blots were washed 3 x 15 min in 0.1% TBST and then incubated with 1:10,000 Rb/Ms-IRDye 680RD secondary antibodies in 0.1% TBST for 1h at room temperature. After 3 x 15 min washes with 0.1% TBST, the bands were detected by the OdysseyCLx Imaging System (LI-COR Biosciences) and quantitatively analyzed by ImageJ software (NIH). Each sample was normalized to its β-actin, then normalized with the corresponding control.

### Patch-clamp recordings Acute slice preparations

Electrophysiology was performed in slices prepared from 2–5 months *Scn2a^gt/gt^* and WT littermates. Mice were deeply anesthetized with ketamine/xylazine (100/10 mg/kg, i.p., 0.1 mL per 10 g of body weight), transcardially perfused, and decapitated to dissect brains into ice-cold slicing solution containing the following (in mM): 110 choline chloride, 2.5 KCl, 1.25 NaH_2_PO_4_, 25 NaHCO_3_, 0.5 CaCl_2_, 7 MgCl_2_, 25 glucose, 1 sodium ascorbate, 3.1 sodium pyruvate (bubbled with 95% O_2_ and 5% CO_2_, pH 7.4, 305–315 mOsm). Acute coronal slices containing frontal cortex and striatum (300-μm in thickness) were cut by using a vibratome (Leica VT1200 S, Germany), and incubated in the same solution for 10 min at 33°C. Then, slices were transferred to normal artificial cerebrospinal fluid (aCSF) (in mM): 125 NaCl, 2.5 KCl, 2 CaCl_2_, 2 MgCl_2_, 25 NaHCO_3_, 1.25 NaH_2_PO_4_, 10 glucose (bubbled with 95% O_2_ and 5% CO_2_, pH 7.4, 305–315 mOsm) at 33°C for 10–20 min and at room temperature for at least 30 min before use. Slices were visualized under IR-DIC (infrared-differential interference contrast) using a BX-51WI microscope (Olympus) with an IR-2000 camera (Dage-MTI).

### Ex vivo electrophysiological whole-cell recordings

All somatic whole-cell patch-clamp recordings were performed from identified striatal MSNs or cortical layer II/III pyramidal neurons. The selection criteria for MSNs were based on morphological characteristics with medium-sized cell bodies presenting polygon or diamond viewed with a microscope equipped with IR-DIC optics (BX-51WI, Olympus), and numerous dendritic spines and their hyperpolarized RMP (lower than –80 mV) based on published method (94). Layer II/III pyramidal cells with a prominent apical dendrite were visually identified mainly by location, shape, and pClampex online membrane test parameters (95).

For whole-cell current-clamp recordings, the internal solution contained (in mM): 122 KMeSO_4_, 4 KCl, 2 MgCl_2_, 0.2 EGTA, 10 HEPES, 4 Na_2_ATP, 0.3 Tris-GTP, 14 Tris-phosphocreatine, adjusted to pH 7.25 with KOH, 295–305 mOsm.

The input resistance (R_input_) was calculated with the equation:

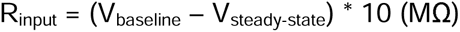

Where V_baseline_ is the resting membrane potential or –80 mV, and V_steady-state_ (V_ss_) is the voltage recorded at 0–10 ms before the end of the –100 pA stimulus.

The RMP, AP threshold, amplitude, fast afterhyperpolarization (AHP), and half-width values were obtained in response to a 20 ms current step of the smallest current to obtain an intact AP, each sweep duration of 1.5 s and start-to-start intervals of 10 s with cells held at the normal RMP or a fixed potential of –80 mV. The RMP, AP threshold, amplitude, AHP, and half-width values were analyzed using the Clampfit 11.4 inbuilt statistics measurements program (Criteria included the baseline, peak amplitude, antipeak amplitude, and half-width). The threshold was defined as the Vm when dV/dt measurements first exceeded 15 V/s.

We used thin-wall borosilicate pipettes (BF150-110-10) with open-tip resistances of 3–5 MΩ. Recordings were performed with an Axon MultiClamp 700B amplifier (Molecular Devices), and data were acquired using pClamp 11.1 software at the normal RMP or a fixed potential of - 80 mV, filtered at 2 kHz and sampling rate at 50 kHz with an Axon Digidata 1550B plus HumSilencer digitizer (Molecular Devices). Slices were maintained under continuous perfusion of aCSF at 32–33°C with a 2–3 mL/min flow. In the whole-cell configuration, recordings with stable series resistance (Rs) 15–30 MΩ were used, and recordings with unstable Rs or a change of Rs > 20% were aborted.

For cell labeling, the internal solution contains 0.1%–0.2% (w/v) neurobiotin tracer. At the end of the electrophysiological recording (about 30 min), slices were treated as previously described (96). Briefly, sections were fixed in 4% paraformaldehyde in 0.1M phosphate buffer (pH 7.4) for 20–30 min at room temperature and subsequently washed 3–4 times for 30 min in 0.1 M phosphate-buffered saline (PBS, pH 7.4) at 4°C. Sections were then incubated in Alexa 488-conjugated streptavidin (overnight at 4C, 1:250 in blocking solution) to visualize neurobiotin.

### hiPSC Lines and Organoid Generation

Detailed methods and reagents are provided in our previous study (97) and the supplemental table. In brief, human induced pluripotent stem cell (hiPSC) lines carrying the *SCN2A* protein-truncating mutation c.2877C>A (p.Cys959Ter) were generated via CRISPR/Cas9 editing. Each genotype has three hiPSC lines. hiPSC colonies were cultured on Matrigel in StemFlex medium. hiPSCs were dissociated with Accutase and seeded in ultra-low attachment 96-well plates with Essential 8 medium supplemented with 10 µM Y27632. After centrifugation at 100 g for 3 min, plates were incubated at 37°C with 5% CO_₂_. At 24 h, media was replaced with Essential 6 containing 2.5 µM dorsomorphin, 10 µM SB-431542, and 1.25 µM XAV-939 for 5 days for neuronal induction via DUAL-SMAD method. On day 6, organoids were transferred to ultra-low attachment 6-well plates in neural induction medium consisting of Neurobasal-A, B-27 without vitamin A, GlutaMAX, and 1:100 penicillin-streptomycin, supplemented with 20 ng/mL FGF2 and 20 ng/mL EGF. From day 22 onward, cerebral organoids were differentiated using 20 ng/mL BDNF, 20 ng/mL NT-3, 200 µM ascorbic acid, 50 µM dibutyryl-cAMP, and 10 µM DHA. Organoids were maintained from day 46 in neural medium supplemented with B-27 Plus with no growth factors until day 150 with media changes every 4–5 days.

### Microelectrode Array (MEA) Recordings from 2D Neuronal Cultures Derived from Human Cerebral Organoids

Detailed methods and reagents are provided in our previous study (97) and the supplemental table. In brief, 3–5 mature (>110 days) organoids were randomly dissociated with 5LJmL of papain-DNase solution and incubated at 37°C with 5% CO_2_ with shaking (80 rpm) for 30LJmin. Single-cell suspension was achieved through mechanical trituration with a flame-polished glass pipette. The supernatant was mixed with inhibitor solution, centrifuged at 300 g for 7LJmins, resuspended in pre-warmed Neurobasal medium, and filtered through a 40 μm mesh.

For MEA recording, ∼7 × 10^4^ cells per well were seeded into a 48-well Cytoview MEA plate pre-coated with 0.1 mg/mL poly-L-ornithine and 10 μg/mL laminin. Cells were cultured in Neurobasal medium supplemented with B-27 without vitamin A, GlutaMAX, and penicillin-streptomycin. From day 7 post-seeding, cultures were switched to BrainPhys medium supplemented with B-27 Plus. Viral transduction was performed on day 7 by adding either control virus (AAV9/TOCitPCamTA, 1.0 × 10^12^ GC/mL) or hK_V_1.1 virus (AAV9/TOKV1CamTA, 1.0 × 10^12^ GC/mL) 1 µL per well. Doxycycline (1 µg/mL) was administered for 7 days to induce expression, followed by a 3-week activation phase without doxycycline. MEA recordings were then conducted Maestro MEA platform (Axion Biosystems). Each well was recorded for 300LJs using AxIS software. Spikes were defined as voltage deflections exceeding 6 standard deviations from the baseline noise. Electrodes registering more than 5 spikes per minute were considered active. Quality control steps include monitoring spike waveform integrity and excluding wells with uneven cell distribution or low viability.

### RT-qPCR

Total RNA was extracted from mouse brains or organoids using the RNeasy Mini Kit (QIAGEN, #74104) following the protocol by the manufacturer. RNA integrity and concentration were assessed using a NanoDrop spectrophotometer. RNA was reverse transcribed to cDNA using the Maxima First Strand cDNA Synthesis Kit (Thermo Fisher Scientific, K1672). Converted cDNAs and corresponding primer sets were combined with Toyobo Thunderbird SYBR qPCR Mix and added in triplets in 96-well plate for quantitative analysis.

The result is read on a C1000 Touch PCR thermal cycler (Bio-Rad). Gapdh and β-actin mRNA levels were used as an endogenous control for normalization using the ΔCt method. In brief, test (T): ΔCt = [Ct (target gene) - Ct (internal control)]; Amount of the target = 2^−ΔCt^.

### Statistics

A set of normality, equal variance, and outlier tests were performed by GraphPad Prism 10 to guide our selection of the most appropriate tests. For comparison between two groups of independent continuous data, if the normality test was significant, the Mann-Whitney U-test (non-parametric) was used; otherwise, the two-tailed unpaired Student’s t-test (parametric) was used. For before-after data from the same animals, two-tailed paired t-test (for two groups) and matched two-way ANOVA (for three or more groups) were used. For independent continuous data with more than two groups, unmatched two-way ANOVA with Tukey correction (parametric) or Kruskal-Wallis with Dunn’s multi-comparison correction (non-parametric) were used.

*Post hoc* multiple comparisons were carried out only when the primary tests showed statistical significance. All data were expressed as mean ± SEM, with a confidence level of 95% (α = 0.05). Specifically, p > 0.05 is indicated as n.s. (no significance), p < 0.05 is indicated as one asterisk (*), p < 0.01 is indicated as two asterisks (**), p < 0.001 is indicated as three asterisks (***), and p < 0.0001 is indicated as four asterisks (****) in all figures. Randomization and blindness were conducted whenever possible to average out the individual differences between litters, housings, body weights, sexes, etc.

## Resource availability

### Lead contact

Further information and requests for resources and reagents should be directed to and will be fulfilled by the lead contact, Yang Yang.

### Materials availability

*Scn2a* gene-trap mice, AAV9/TOKv1CamTA, AAV9/NegCTRLCam, and AAV9/TOCitPCamTA are generated and used in this study.

### Data and code availability

All data used in this study are reported in the Supplementary Materials. Any additional information required to reanalyze the data reported in this paper is available from the lead Contact upon request.

## Author contributions

Z.Z., J.Z., X.C., and Y.Y. designed research; Z.Z., J.Z., X.C., B.D., P.J.S., M.S.H., A.D.A. and M.T.T. performed experiments; Y.V., R.P.G., E.P.R., provided unpublished reagents; S.K., P.M., H.K., M.J.R., Y.Z., C.Y., N.A.L., D.W., G.G., and R.S. participated in research design/data analysis; Z.Z., J.Z., and Y.Y. wrote the manuscript with inputs from all authors.

## Acknowledgments

We sincerely thank Dr. Steve C. Danzer, Dr. Danielle Tapp, and Dr. Kimberly Kraus from Cincinnati Children’s Hospital Medical Center for their invaluable guidance in EEG recording. The research reported in this publication was supported by the NINDS of the NIH (R01NS117585 and R01NS123154 to Y.Y.; and NS097726 to E.P.R). X.C. was supported by the AES Postdoctoral Research Fellowship. The authors gratefully acknowledge support from the Purdue Institute for Drug Discovery and the Purdue Institute for Integrative Neuroscience for additional funding support. Yang Lab is grateful to the *FamilieSCN2A* Foundation for the Hodgkin-Huxley Research Award to Y.Y. and the Action Potential Grant support to X.C. and J.Z. This project was supported in part by the Indiana Spinal Cord & Brain Injury Research Fund and the Indiana CTSI, funded in part by UL1TR002529 from the NIH. The Yang lab appreciates the bioinformatics support of the Collaborative Core for Cancer Bioinformatics (C3B) with support from the Indiana University Simon Comprehensive Cancer Center (Grant P30CA082709), Purdue Institute for Cancer Research (Grant P30CA023168), and the Walther Cancer Foundation. The content is solely the responsibility of the authors and does not necessarily represent the official views of the Indiana State Department of Health and the National Institutes of Health.

## Declaration of AI-assisted technologies in the writing process

ChatGPT 4.5 was used in this manuscript to improve grammatical accuracy, language fluency, and readability. We ensured that the intended meaning of the sentences remained strictly unchanged. The authors carefully reviewed and further edited each sentence to guarantee they complied with scientific rigor. AI-assisted tools were not used in any images or other multimedia. The authors take full responsibility for the content of the publication.

## Supplemental Figures

**Supplemental Figure 1.**
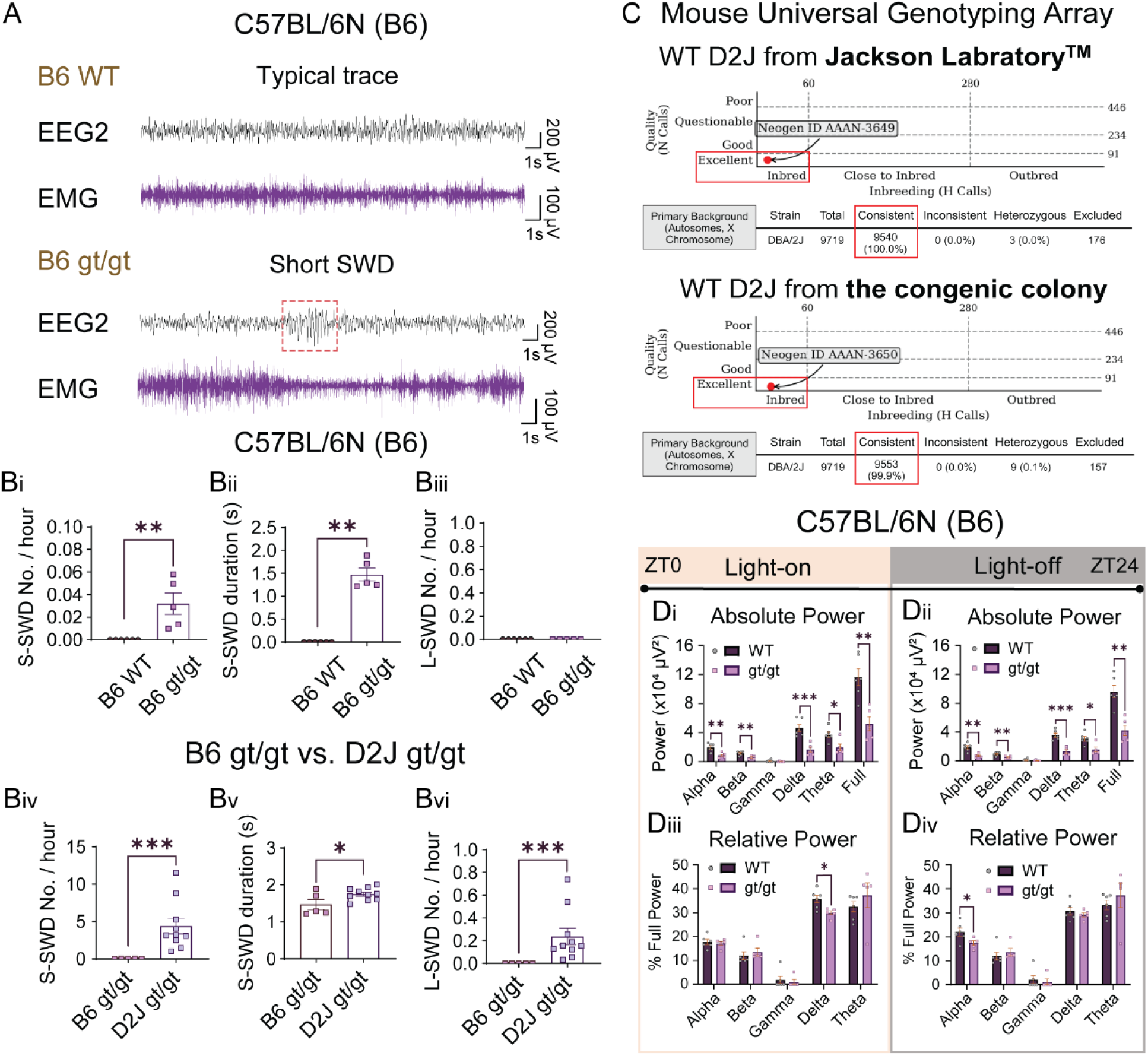
C57BL/6N mice with severe *Scn2a* deficiency show weak spike-wave discharges and power spectra alterations. (A) Representative EEG traces showing the complete absence of spike-wave discharges in the B6 WT and a small, short SWD (red dashed box) in the B6 *Scn2a^gt/gt^* mice. (Bi-iii) Quantification of SWDs in the B6 WT and B6 *Scn2a^gt/gt^* mice. There was a significant increase in SWD frequency and duration in the B6 *Scn2a^gt/gt^* compared to the B6 WT mice, which had no SWD detected. Additionally, long SWDs with longer than 3.5 s duration were negligible in the B6 *Scn2a^gt/gt^*. (Biv-vi) Comparison of SWD quantifications between B6 *Scn2a^gt/gt^*and D2J *Scn2a^gt/gt^*. Although B6 *Scn2a^gt/gt^* showed a significant SWD increase compared to B6 WT, its frequency and duration scale was much less compared to the D2J *Scn2a^gt/gt^*. D2J *Scn2a^gt/gt^*also showed long absence seizures which were negligible in the B6 *Scn2a^gt/gt^*. (C) Giga mouse universal genotyping array (GigaMUGA) showed that mice from the D2J congenic colony had 99.9% genome consistency with pure wildtype D2J purchased from Jackson Laboratory. (D) Quantification of 1-week EEG2 power spectra during the light-on and light-off period for the B6 WT and *Scn2a^gt/gt^* mice. Similar to the D2J, B6 *Scn2a^gt/gt^* mice showed an overall reduction of absolute power. The % delta power and % alpha power were also decreased during the light-on and light-off period, respectively. Data are presented as mean ± SEM. Statistical analyses: Non-parametric Mann-Whitney U test (Bi-Biv)(Bvi). Unpaired Student’s t test (Bv)(Di-Div). *p < 0.05; **p < 0.01; ***p < 0.001; ****p < 0.0001. Exact p values can be found in Supplemental Table 1.

**Supplemental Figure 2.**
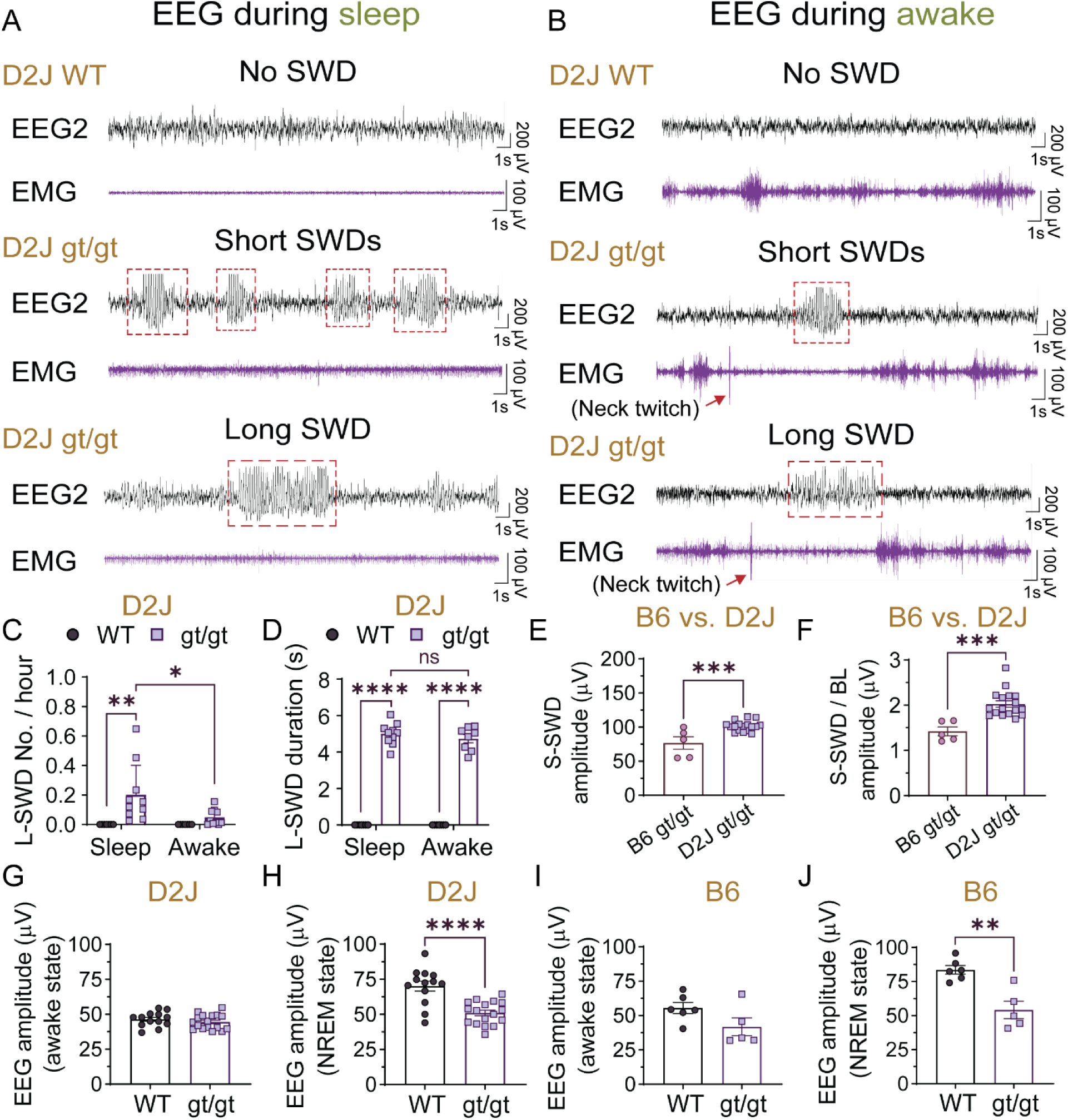
The majority of the long SWDs occur during the sleep state instead of the awake state in the D2J *Scn2a^gt/gt^* mice. (A) Representative EEG traces of D2J WT and D2J *Scn2a^gt/gt^* mice showing typical epileptiform events during the sleep (i.e., NREM) stage. (B) Representative EEG traces of D2J WT and D2J *Scn2a^gt/gt^*mice showing typical epileptiform events during the awake stage. Note the characteristic ‘neck twitch’ behavior which often happens at the onset of a long SWD event. (C) and (D) Quantification of the frequency and duration of absence seizures of D2J WT and D2J *Scn2a^gt/gt^* mice during the sleep vs. awake state. There were significantly fewer long SWDs in D2J *Scn2a^gt/gt^* during the awake state compared to sleep. The duration of L-SWDs was similar in both states. (E) and (F) Quantification of the short SWDs (S-SWD) absolute and relative (i.e. to BL) EEG amplitude (μV) in B6 vs. D2J *Scn2a^gt/gt^* mice. D2J *Scn2a^gt/gt^*mice have significantly higher S-SWD amplitude compared to the B6 *Scn2a^gt/gt^*mice. (G–J) In both B6 and D2J strains, *Scn2a^gt/gt^* mice show a significant reduction of baseline EEG amplitude during the deep sleep (i.e. NREM) state and no difference during the awake state. Data are presented as mean ± SEM. Statistical analyses: Non-parametric Mann-Whitney U test (S2C)(S2D). *p < 0.05; **p < 0.01; ***p < 0.001; ****p < 0.0001. Exact p values can be found in Supplemental Table 1.

**Supplemental Figure 3.**
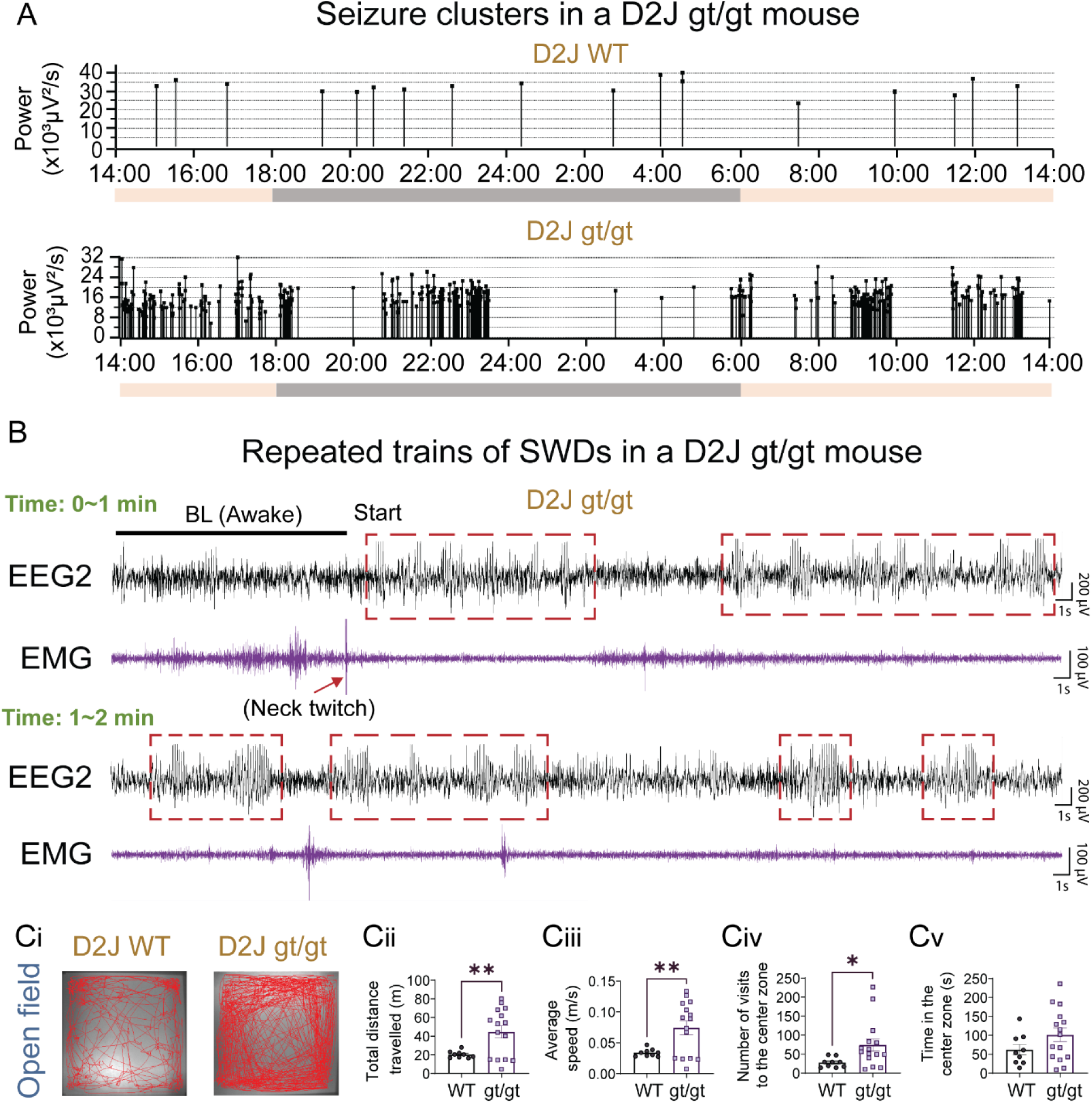
D2J *Scn2a^gt/gt^*mice show SWD clusters in EEG and hyperactivities in the open-field test. (A) Representative images show that the high frequency of SWDs in D2J *Scn2a^gt/gt^* mice occurred during specific time windows over a 24-hour period. In comparison, the D2J WT mice have much fewer SWDs, which tend to distribute randomly throughout the 24 hours. The beige color indicates a light-on period, and the grey color indicates a light-off period. (B) EEG2 and EMG recording of a D2J *Scn2a^gt/gt^* mouse showing an example of repeated trains of SWDs which lasted around 2 minutes. Note the characteristic ‘neck twitch’ behavior from the EMG recording at the beginning of this long epileptiform event. Red dashed boxes show many consecutive repeated mature and immature SWDs in this mouse. (C) Representative tracking plots and quantifications indicate that D2J *Scn2a^gt/gt^* mice traveled significantly greater distances and at higher speeds in the open field test. D2J *Scn2a^gt/gt^*mice demonstrated significantly more crossings in the center zone, yet there was no increase in the time spent in the center zone. Data are presented as mean ± SEM. Statistical analyses: Non-parametric Mann-Whitney U test (Cii-Cv). *p < 0.05; **p < 0.01; ***p < 0.001; ****p < 0.0001. Exact p values can be found in Supplemental Table 1.

**Supplemental Figure 4.**
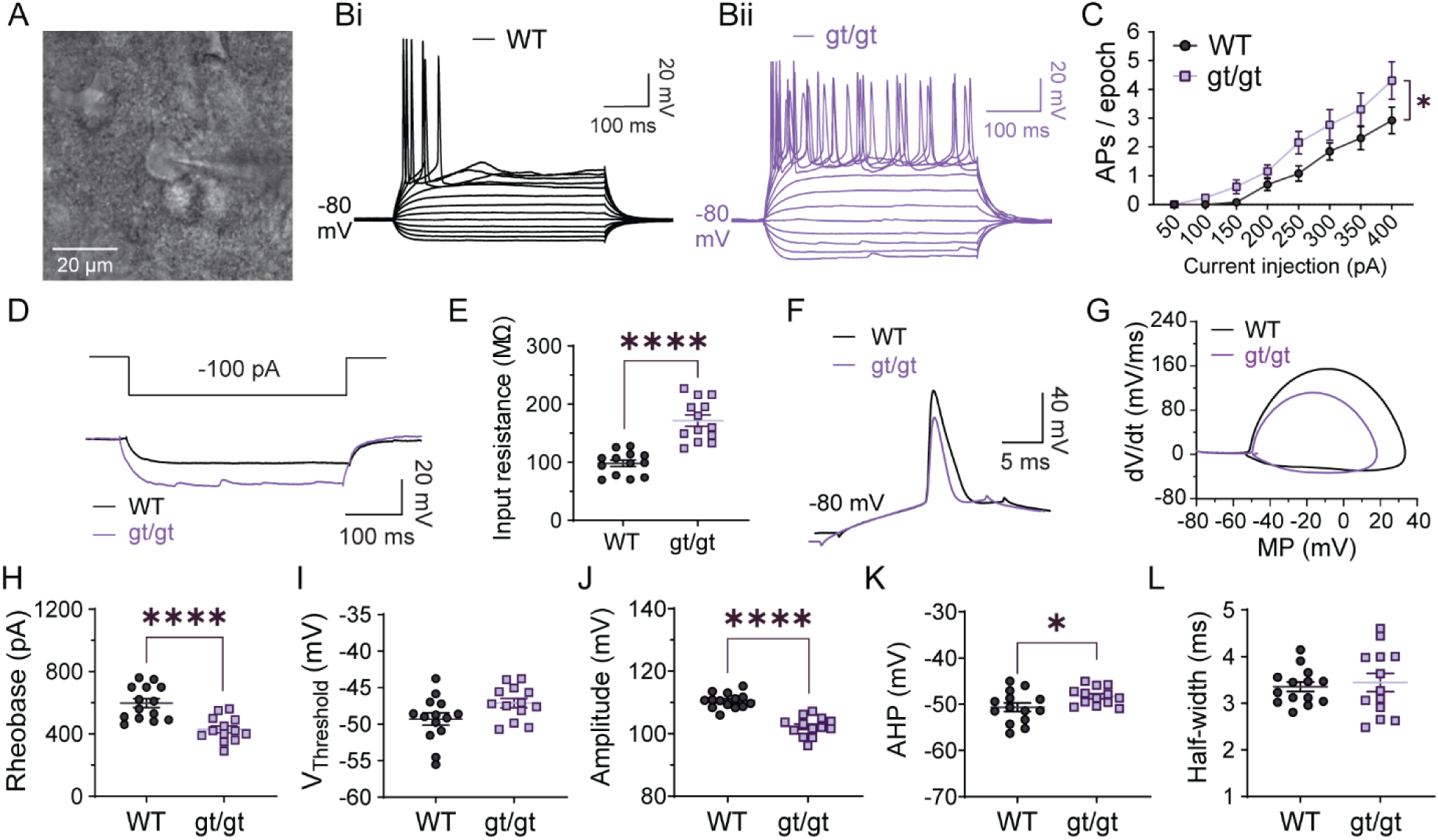
Layer 2/3 pyramidal neurons in the somatosensory cortex of D2J *Scn2a^gt/gt^* mice show intrinsic hyperexcitability when holding at –80 mV. (A) A representative image showing a pyramidal cell at the layer 2/3 of the somatosensory cortex (SSC) being patched. (B-C) Representative traces and quantifications show a trend of increased AP firing of D2J *Scn2a^gt/gt^* mice in response to a step increase in current injection. (D) Representative traces in response to –100 pA injection. (E) Layer 2/3 pyramidal neurons in the D2J *Scn2a^gt/gt^* mice had significant increases in input resistance when holding at –80 mV. (F-L) Layer 2/3 pyramidal neurons in the D2J *Scn2a^gt/gt^*mice had different AP shapes, decreased rheobase, AP amplitude, and AP half-width compared to the D2J WT mice. The voltage threshold and fast after-hyperpolarization (AHP) were unchanged. Data are presented as mean ± SEM. Statistical analysis: Two-way matched ANOVA (C); Unpaired Student’s t test (E)(H-L). *p < 0.05; **p < 0.01; ***p < 0.001; ****p < 0.0001. Exact p values can be found in Supplemental Table 1.

**Supplemental Figure 5.**
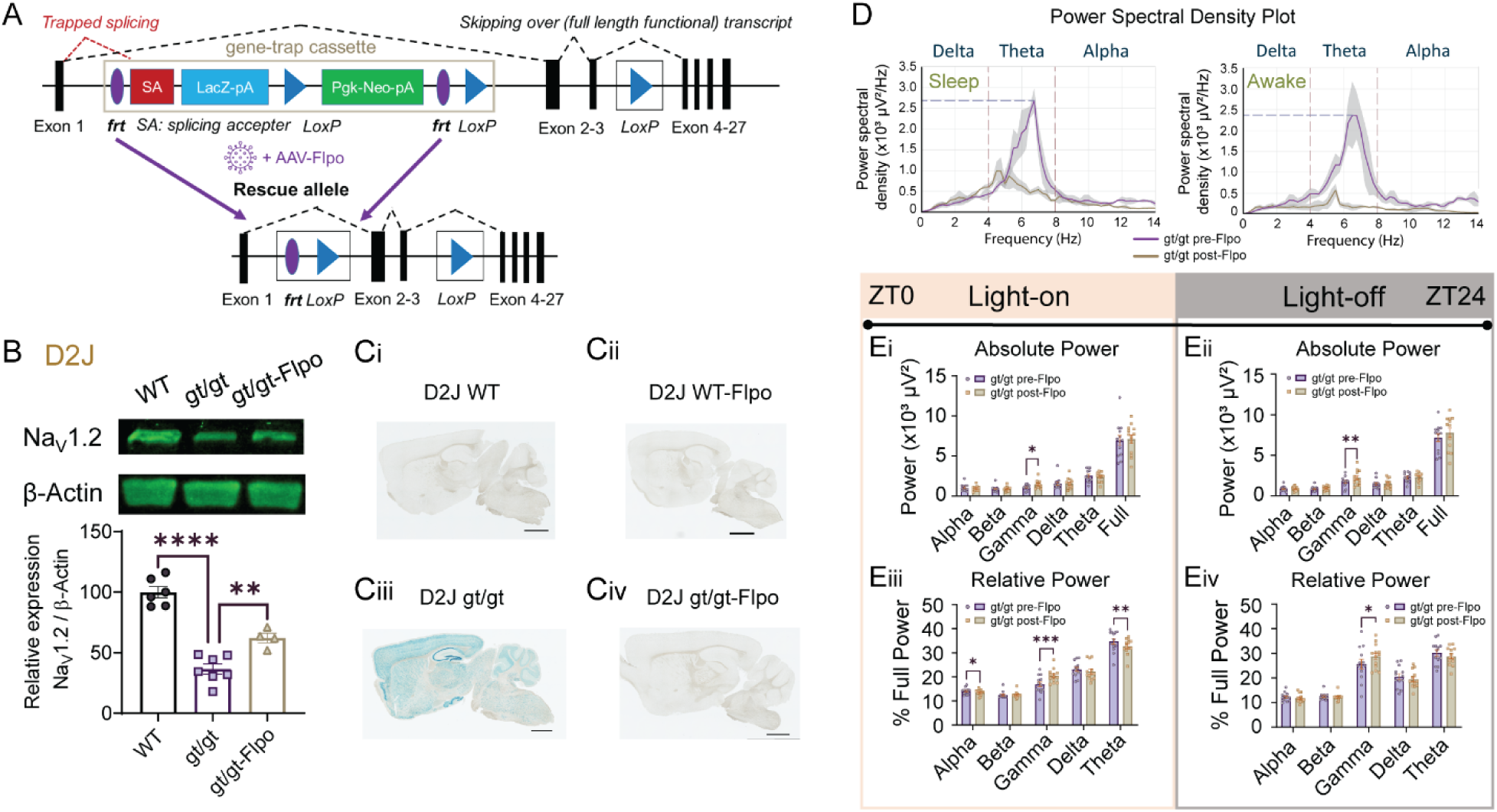
β-galactose staining and western blot results showed that systemic AAV-PHP.eB-Flpo injection at the adult state removed LacZ expression and partially rescued Na_V_1.2 expression. (A) A schematic of the gene-trap constructs and rescue methods. (B) Western blot showing partial Na_V_1.2 protein level rescue after adult tail-vein AAV-Flpo injection to remove the gene-trap cassette. (C) β-galactose staining demonstrates the removal of the gene-trap cassette, which contains the *LacZ* domain. (D) Representative power spectral density plots show that D2J *Scn2a^gt/gt^* mice had significantly higher power density in the theta band pre-Flpo injection (violet) compared to post-Flpo injection (brown). (E) Quantification of 1-week EEG recording showed that D2J *Scn2a^gt/gt^* had a significant increase in absolute and relative gamma power post-Flpo injection. During the light-on period, D2J *Scn2a^gt/gt^* had a significant decrease in relative alpha power and theta power after Flpo injection. Data are presented as mean ± SEM. Statistical analyses: Two-tailed Pearson correlation coefficients (E-H). *p < 0.05; **p < 0.01; ***p < 0.001; ****p < 0.0001. Exact p values can be found in Supplemental Table 1.

**Supplemental Figure 6.**
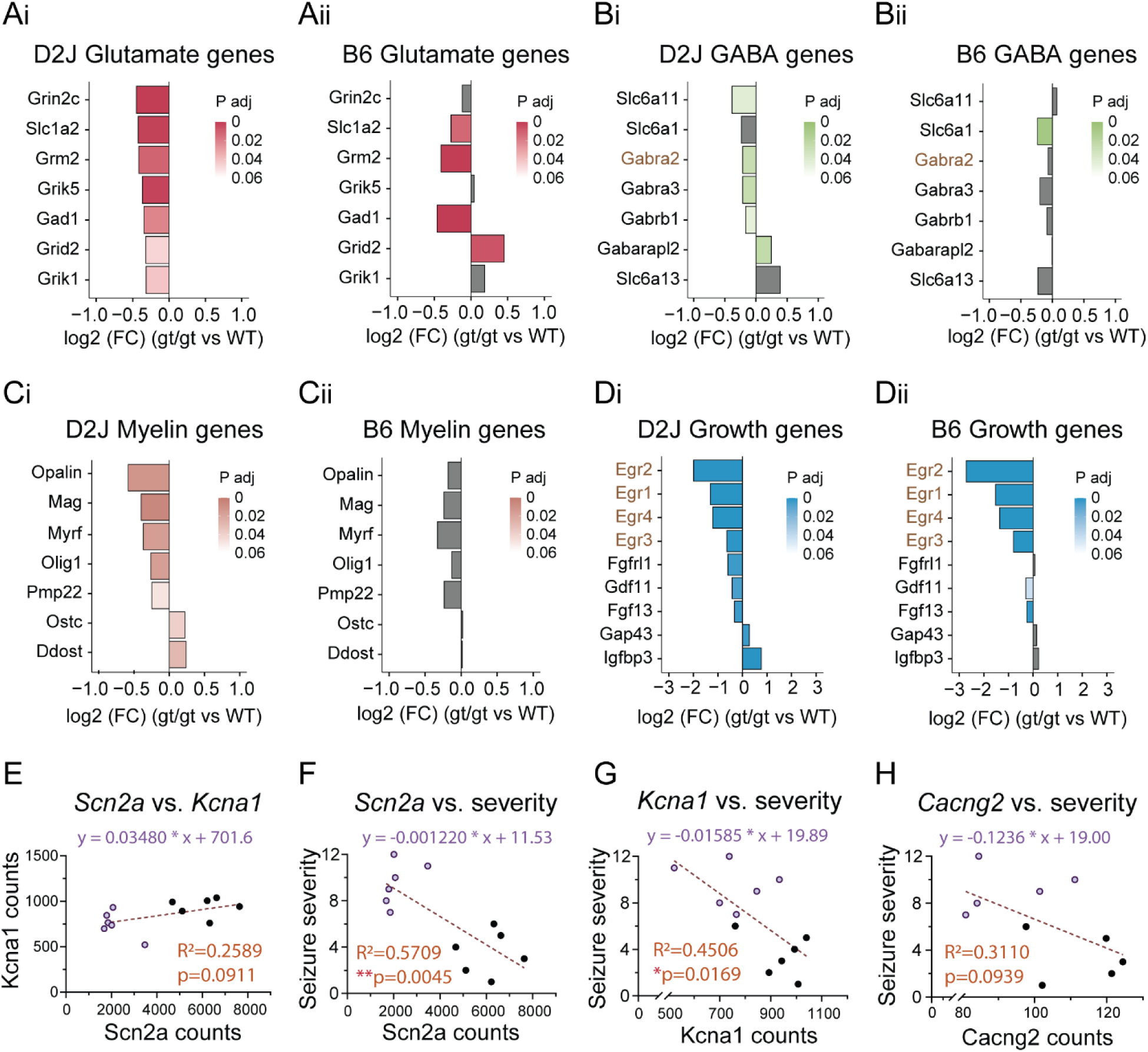
Glutamate, GABA, myelin, and growth-related genes were differentially altered in D2J vs. B6 *Scn2a^gt/gt^*mice. (A-B) Side-by-side comparison of glutamate and GABA-related genes significantly downregulated in the D2J *Scn2a^gt/gt^* and B6 *Scn2a^gt/gt^* mice relative to corresponding WTs. Most of these genes were differentially regulated except for *Slc1a2*, *Grm2*, and *Gad1*, which were consistently downregulated in both D2J and B6 gt/gt mice. (C) A set of myelin-related genes significantly down/upregulated in the D2J *Scn2a^gt/gt^*mice were non-significant in the B6 *Scn2a^gt/gt^* mice. (D) A set of growth-related genes (esp. *Egr* family) were consistently down/upregulated in both D2J and B6 *Scn2a^gt/gt^* mice. (E) There was no correlation between *Scn2a* and *Kcna1* gene expression based on normalized count. (F-H) Absence seizure severity in the D2J *Scn2a^gt/gt^* mice was significantly correlated with *Scn2a* and *Kcna1* gene expression but not with *Cacng2* expression. Data are presented as mean ± SEM. Statistical analyses: Two-tailed Pearson correlation coefficients (E-H). *p < 0.05; **p < 0.01; ***p < 0.001; ****p < 0.0001. Exact p values can be found in Supplemental Table 1.

**Supplemental Figure 7.**
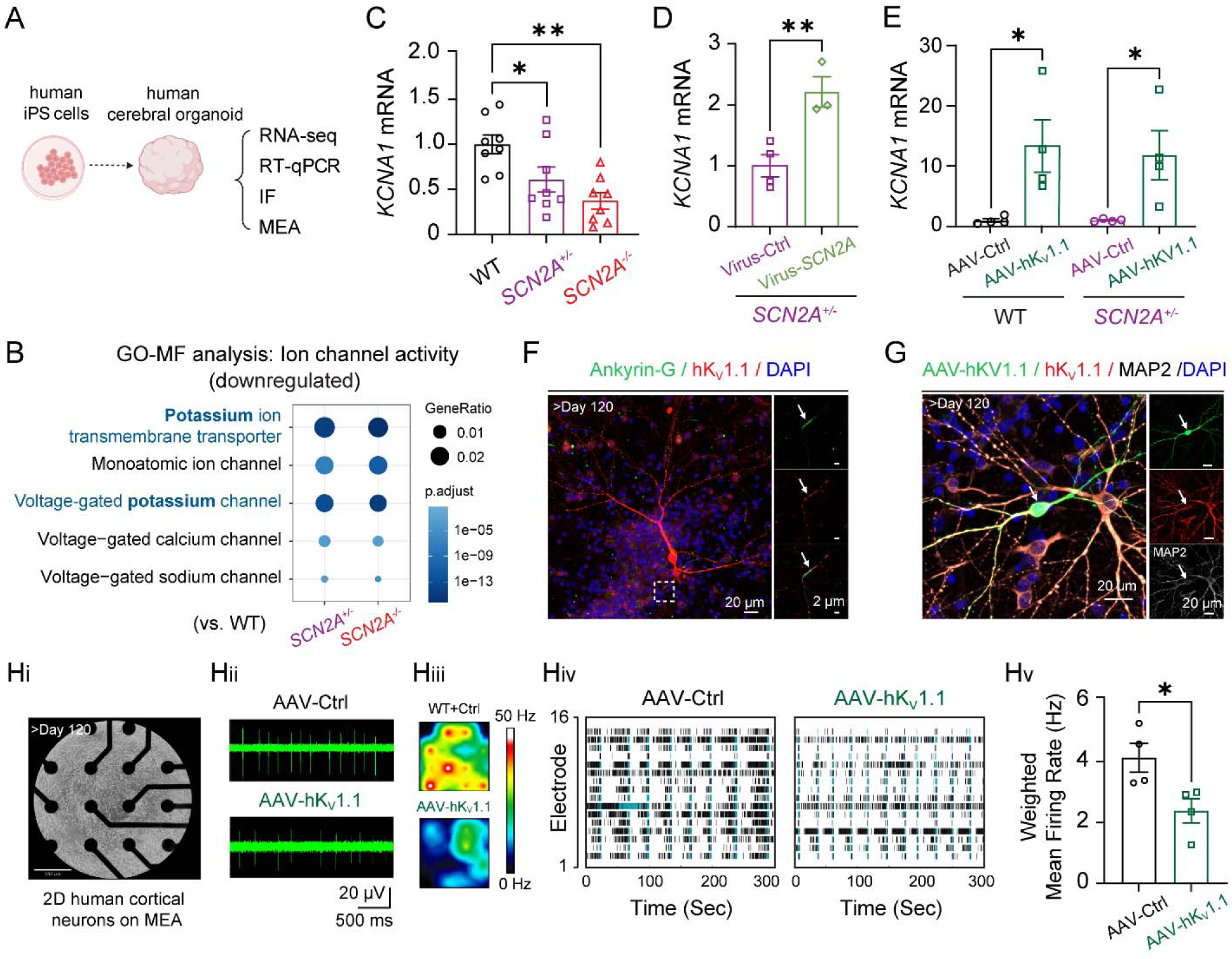
K*C*NA1 is downregulated in human brain organoids with *SCN2A* deficiency. (A) Schematic of the human cerebral organoid experimental design. *SCN2A* hiPSC lines carrying protein-truncating variant (*SCN2A^+/-^*and *SCN2A^−/-^*) were generated via CRISPR/Cas9 genome editing. IF: immunofluorescence; MEA: multielectrode array. (B) Gene ontology (GO) molecular function analysis from bulk-RNA sequencing indicates significant downregulation of genes associated with voltage-gated potassium channel (K_V_) in *SCN2A*-deficient organoids. (C-E) RT-qPCR analyses demonstrate a significantly reduced *KCNA1* mRNA level in *SCN2A*-deficient organoids compared with WT controls (C), which was reversible by viral vector-mediated *SCN2A* restoration (D). Notably, AAV-hK_V_1.1 successfully elevated *KCNA1* expression in brain organoids (E). (F-G) Immunofluorescence imaging indicates localization of K_V_1.1 at the axon initial segment (AIS, marked by Ankyrin-G), in brain organoids. Scale bars: 20 µm (main), 2 µm (insets). AAV-hK_V_1.1 effectively transduces human neurons in organoids (>120 days) (G). Scale bars: 20 µm. (Hi-Hv) Multielectrode array recordings reveal that exogenous hK_V_1.1 expression reduces neuronal action potential firing in dissociated human neurons. Data are presented as mean ± SEM. Statistical analyses: One-way ANOVA and Student’s t test: *p < 0.05; **p < 0.01; ***p < 0.001; ****p < 0.0001. Exact p values can be found in Supplemental Table 1.

